# Delayed aneuploidy stress response of neural stem cells impairs adult lifespan in flies

**DOI:** 10.1101/392746

**Authors:** Mihailo Mirkovic, Leonardo G. Guilgur, Diogo Passagem-Santos, Raquel A. Oliveira

## Abstract

**Summary:** Studying aneuploidy during organism development has strong limitations, as chronic mitotic perturbations used to generate aneuploidy result in lethality. We developed a genetic tool to induce aneuploidy in an acute and time controlled manner during *Drosophila* development. This is achieved by reversible depletion of cohesin, a key molecule controlling mitotic fidelity.

Larvae challenged with aneuploidy hatch into adults with severe motor defects shortening their lifespan. Neural stem cells, despite being aneuploid, display a delayed stress response and continue proliferating, resulting in the rapid appearance of chromosomal instability, complex array of karyotypes and cellular abnormalities. Notably, when other brain cell-lineages are forced to self-renew, aneuploidy-associated stress response is significantly delayed, indicating that stemness state confers resistance to aneuploidy. Sparing solely the developing brain from induced aneuploidy is sufficient to rescue motor defects and adult lifespan, suggesting that neural tissue is the most ill-equipped to deal with developmental aneuploidy.

**Highlights:**

- Reversible depletion of cohesin results in just a round or two of aberrant cell divisions, generating high levels of aneuploidy.
- Larvae challenged with aneuploidy during development hatch into impaired adults.
- Few cell cycles are sufficient for chromosomal instability to emerge from a previously stable aneuploid state.
- Neural stemness delays aneuploidy stress response.
- Protecting only the neural tissue from aneuploidy rescues adult abnormalities and lifespan.

## Introduction

Aneuploidy, a state of chromosome imbalance, was observed over a century ago by Theodor Boveri. Since then, numerous studies have shown that aneuploidy is largely detrimental both at cellular and organism level. In multicellular organisms chromosome gain or loss results in lethality or developmental defects (1, 2). At the cellular level, studies in yeast and cell culture have demonstrated that aneuploidy has a high fitness cost for the cell, as unbalanced karyotypes lead to activation of multiple stress response pathways, resulting in reduced proliferation, cell cycle arrest, or cell death (Reviewed in (3). The aneuploidy stress response and consequential drop in fitness seems at odds with the hypothesized role of aneuploidy in promoting malignancy, which is usually marked by over-proliferation (4). Ninety percent of solid tumors harbor whole chromosome gains and/or losses (5). Therefore, although usually detrimental to cell fitness, aneuploidy and its effects on cell proliferation can be context dependent, which emphasizes our need for a better understanding of the immediate and ultimate consequences of this abnormal cellular condition in a tissue context and through development.

Study of aneuploidy *in vivo* is challenging since somatic aneuploidy is a rare event and is difficult to capture and trace in real time due to several constraints: i) Cells are equipped with surveillance mechanisms that prevent chromosome mis-segregation (e.g Spindle Assembly Checkpoint (SAC) (Reviewed in (6) making naturally occurring aneuploidy events virtually impossible to evaluate; ii) experimentally-induced aneuploidy, by compromising mitotic fidelity, if often of low prevalence, as it has been demonstrated for several mammalian (7) (8) and *Drosophila* tissues (9, 10) and iii) induction of somatic or constitutional aneuploidy in metazoans relies on chronic mitotic perturbation (Listed in (11) which usually causes embryonic lethality (Reviewed in(12) as a result of progressive accumulation of damage in the developing organism. Thus, from these studies, it is impossible to disentangle short term and long term consequences of aneuploidy, or to examine kinetics of the response to aneuploid state during development. To circumvent these limitations, we generated a genetic system with the power to induce aneuploidy in an acute and time-controlled manner, in all the dividing tissues of the developing *Drosophila*. The tool is based on reversible depletion of cohesin, a key molecule regulating mitotic fidelity (13, 14). Cohesin is a tripartite ring complex, composed by SMC1, SMC3 and the bridging kleisin subunit RAD21 (15, 16). The primary mitotic role of cohesin is to mediate sister chromatid cohesion, by topologically entrapping DNA fibers from neighboring chromatids (17, 18). Cells entering mitosis with premature loss of cohesion and sister chromatid separation activate the Spindle Assembly Checkpoint (SAC) resulting in prolonged mitosis (14, 19). During this SAC-dependent mitotic delay, chromosomes are shuffled from one cell pole to the other by the mitotic spindle (19). Consequently, chromosome shuffling induces genome randomization and aneuploidy upon mitotic exit with a theoretical rate of nearly 100%. Our engineered system enables a quick restoration of this complex shortly after its inactivation, thereby restricting mitotic abnormalities to a short time-frame, concomitantly with the generation of high levels of aneuploidy. Using such tool, we dissect the kinetics of aneuploidy response across various cell/tissue types and developmental timings.

## Results

### A genetic system for acute and time-controlled generation of aneuploidy in a developing organism

To induce aneuploidy in an acute and time-controlled manner, we developed a genetic system based on rapid removal of cohesin complex, the molecular glue that holds sister chromatids together. To prevent a chronic cohesion depletion state and restrict mitotic failure to a single cell cycle, our genetic system is able to induce cohesin inactivation, followed by subsequent cohesion rescue. The system relies on the artificial cleavage of a modified version of the RAD21 cohesin subunit that contains TEV protease cleavage sites (RAD21-TEV). As previously shown, this system is very efficient at inactivating cohesin upon expression of the exogenous Tobacco Etch Virus (TEV) protease, induced by a heat-shock promoter, resulting in long-term inactivation of this complex (>24h) (20). To restrict cohesin impairment, we modified this system by promptly rescuing cohesin integrity through the expression of TEV-resistant RAD21 protein (RAD21-WT) right after TEV-mediated inactivation. For this purpose, RAD21-WT expression is under the control of UAS *promoter (UAS-Rad21-wt-myc)* that is induced by a Gal4 protein induced concomitantly with the TEV protease (also under a heat-shock promoter, *HSprom-GAL4)* (Figure 1A). Given that the TEV protease is under a direct control of heat-shock *promoter*, whereas RAD21-WT relies on a dual expression-system *(Gal4-UAS)*, we anticipated that the temporal delay in RAD21-WT expression relative to the induction of TEV protease would lead to a short time window of cohesin inactivation (RAD21 cleavage) (Figure 1A).

**Figure 1.**
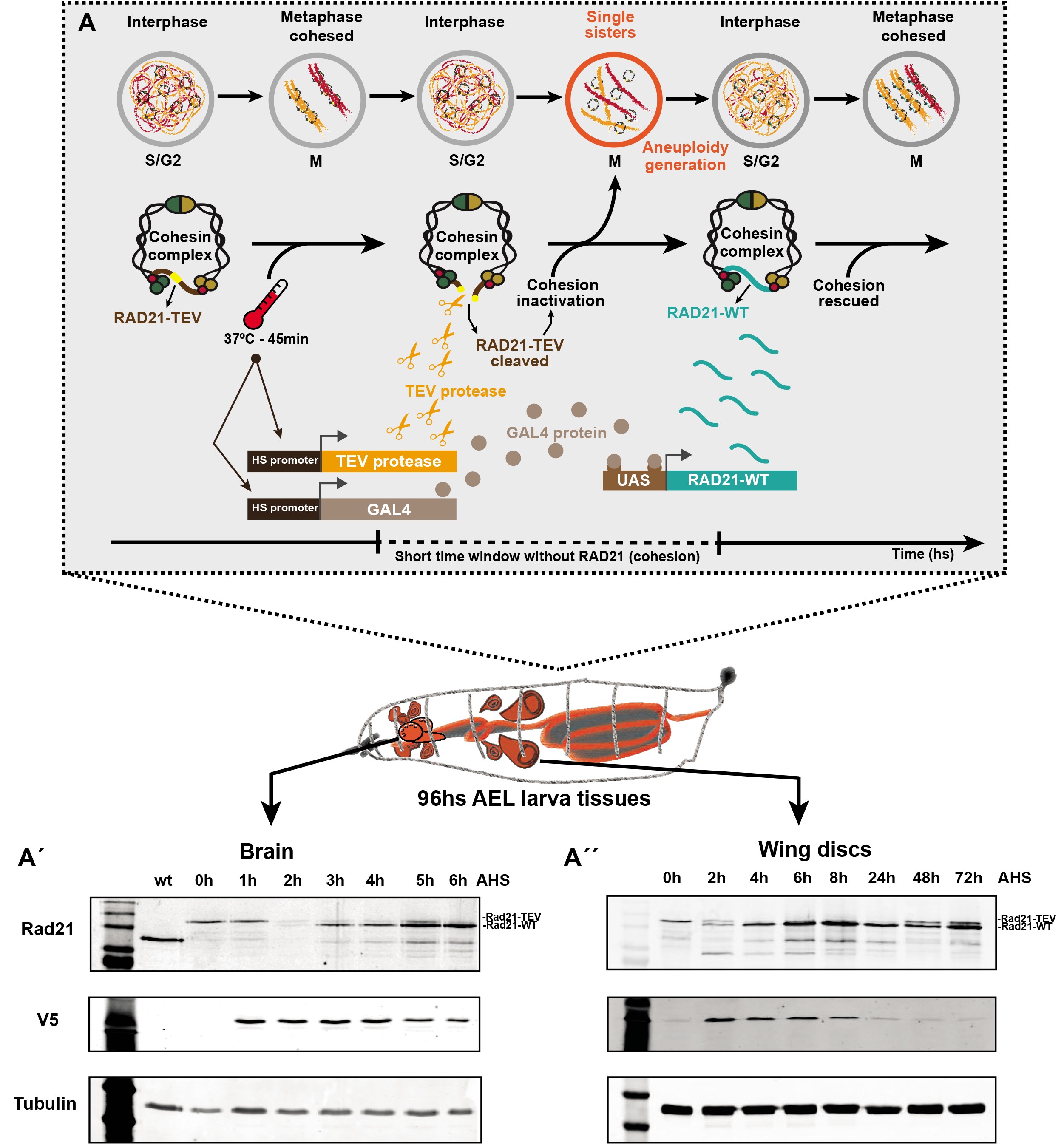
Reversible system for acute cohesin depletion and generation of aneuploidy in the developing *Drosophila*. **A to A’’: A-** The system relies on the cleavage of RAD21 Cohesin subunit (RAD21-TEV) by the heat-shock induced expression of TEV protease. Cohesin cleavage is rapid and the RAD21-TEV is completely depleted within hours after the heat-shock (see western blots A’, A’’, RAD21 upper band). Cells entering into mitosis with loss of cohesion experience premature sister chromatid separation, chromosome shuffling and as consequence generation of aneuploidy daughter cells with a theoretical rate of nearly 100%. To restrict mitotic failure to a single cell cycle, shortly after aneuploidy has been generated, cohesion is rescued by inducing the expression of TEV-resistant RAD21 protein (RAD21-WT). The RAD21-WT subunit is reliant on heat-shock Gal4 protein expression, coupled to the expression of the UAS-RAD21-WT and occurs within hours after the heat-shock (see western blots A’, A’’, RAD21 lower band). **A’and A’’-** Western blots displaying RAD21-TEV cleavage and RAD21-WT restoring cohesin function in the 3^rd^ *instar* larvae brains and wing discs. Tissues samples were taken at different hours after the heat-shock induction (AHS). V5 antibody evidences the TEV protease expression (from *hspr-nlsV5TEV)* and Tubulin antibody was used as loading control.

To test this, we probed for the kinetics of TEV-mediated cleavage of RAD21-TEV and synthesis of RAD21-WT in different tissues of the developing larvae. After heat shock, both *Drosophila* larvae brains and wing discs, showed similar kinetics of the TEV-sensitive RAD21 disappearance followed by the appearance of RAD21-WT (Figures 1A’ and 1A’’). The timing of protein depletion/re-establishment differs slightly among different tissues or developmental stages, but leads on average to a period of ~1 hour without cohesin (Figures 1A’; 1A’’ and S1B).

### Reversible removal of cohesin results in a single round of mitotic abnormalities and consequent aneuploidy

The cohesive function of cohesin is established in S-phase, concomitantly with DNA replication. Once stabilized on the replicated genome, cohesive cohesin complexes do not turn over (21). As such, loss of cohesin using our system will affect sister chromatid cohesion in all cells that are in S/G2/M phase during the short period between TEV protease expression and synthesis of RAD21-WT (Figure 1A). In addition to its canonical cohesive function, cohesin has also been recently implicated in other interphase functions, including regulation of gene expression (22). In contrast to the cohesive pool, these cohesin molecules are known to be highly dynamic (21, 23). Moreover, cohesin-mediated loops were recently reported “memorable” and quickly reformed upon cohesin re-establishment (24). We therefore anticipated that this function should not be severely affected by our system. In sharp contrast, mitotic errors induced upon cohesin cleavage are irreversible as there is no way to restore cellular ploidy after a compromised round of mitosis.

Whereas canonical chronic mitotic perturbations lead to several rounds of mitotic failures, our novel genetic system should lead to cohesion defects only in the first mitosis following the heat-shock, as the expression of RAD21-WT should be able to rescue cohesion in the subsequent cell cycle, if given enough time (Figure 1A). To confirm that our genetic system works as anticipated, we focused our analysis on two different cycling tissues from the larva: the developing brain and the epithelial wing discs.

The developing brain of *Drosophila* is an excellent model to study the consequences of developmental aneuploidy. The well characterized cell lineages of the tissue in combination with our tractable system to induce mis-segregation of chromosomes offer a unique opportunity to trace the fate of aneuploid cells in real time and analyze their effect on the nervous system development. Through larval development ~100 large neural stem cells called Neuroblasts (Nbs) (25) located in the central brain (CB) region divide asymmetrically to self-renew and generate distinct neuronal lineages via differentiating progeny (26).

We evaluated, by live cell imaging, mitotic fidelity in these Nbs using two independent criteria to estimate the state of sister chromatid cohesion: i) the presence of single sisters (a direct consequence of cohesion loss), as opposed to metaphase chromosome alignment and ii) the time cells spend in mitosis, given that premature loss of sister chromatid cohesion is known to activate the SAC and delay mitotic exit (Mirkovic et al., 2015).

As expected, the first division after the heat shock results in full cohesin cleavage in Nbs, followed by cohesin rescue in subsequent divisions (Movies S1 and S2). The fast cell cycle of Nbs, coupled with continued proliferation of these cells despite their abnormal genome content (further discussed below), enables analysis of mitotic fidelity throughout several consecutive divisions in great detail. Consistently, in the first mitosis AHS, 95% of Nbs contain single sisters, and exhibit mitotic delay and chromosome shuffling (Figures 2A and 2B). In the subsequent mitosis, however, normal cohesion is observed in ~80% of the Nbs, with clear metaphases and a shorter mitotic delay (Figures 2A; 2B and 2C). Finally, during the third cell division AHS, the mitotic timing and the cohesive state of Nbs are comparable to heat-shocked controls (Figures 2A; 2B and 2C). Similar results were obtained for larvae heat-shocked at earlier stages of development (Figure S1A to A’’).

**Figure 2.**
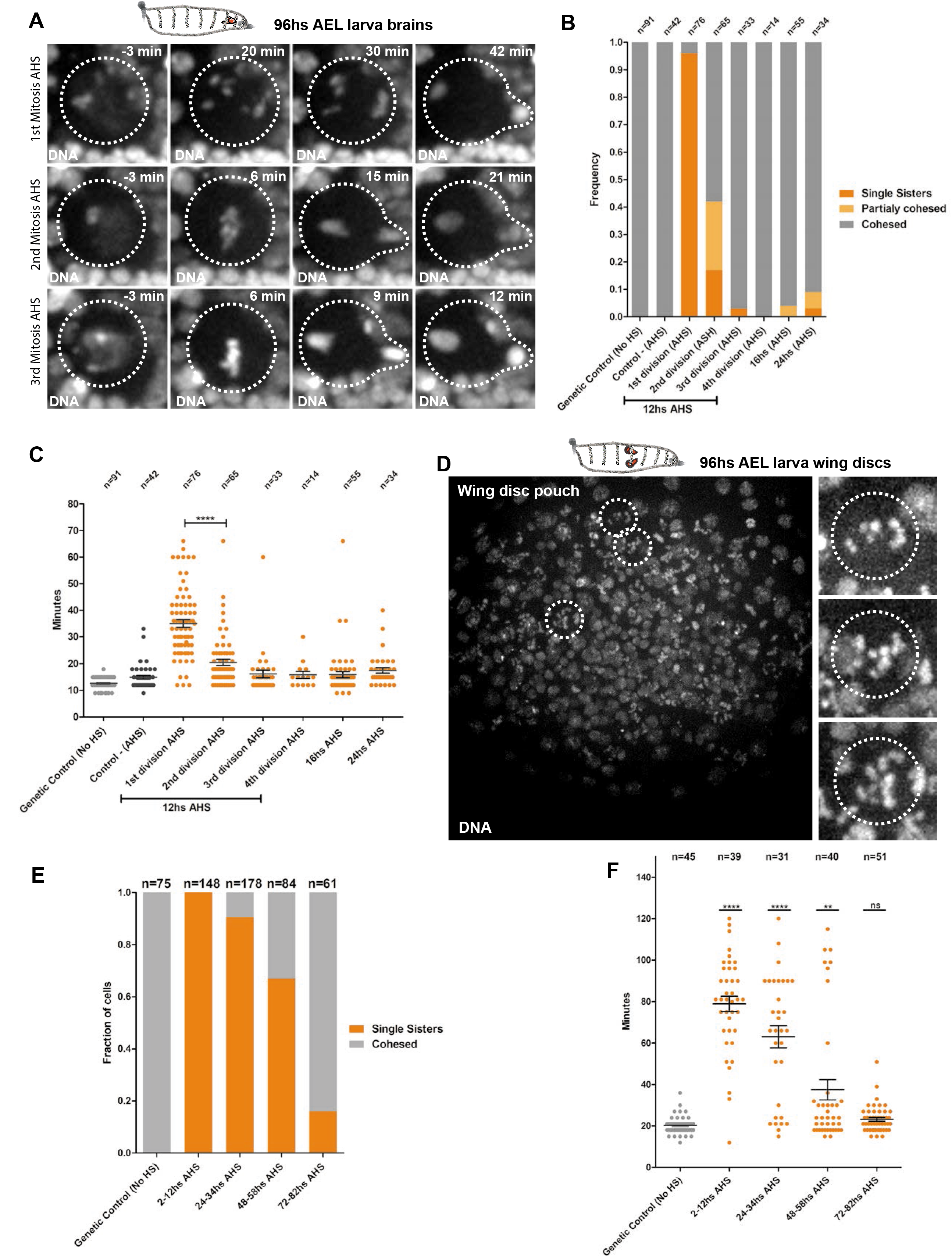
Kinetics of reversible loss of cohesin in the 3^rd^ *instar* brains and the wing discs. **A-** Stills from live imaging of Nbs after the heat-shock (full movies available in supplementary movie 1 and 2). Transient loss of cohesion results in a round of defective mitosis and genome shuffling (1^st^ division AHS). After this round of division, the following mitosis (2^nd^ and 3^rd^ divisions AHS) shows the restoring of cohesion function and mitotic fidelity in larvae Nbs. Dashed circles display the mitotic figures of the dividing Nbs. **B-** Quantification of Cohesive states of 3^rd^ *instar* larvae Nbs following RAD21-TEV cleavage and RAD21-WT restoring of cohesin function. More than 80% of the 2^nd^ divisions AHS are already totally or partially cohesed. **C-** Quantification of mitotic timing and delay caused by Spindle Assembly Checkpoint activation after RAD21-TEV cleavage and RAD21-WT rescue in the 3^rd^ *instar* Nbs (2 to 24hs AHS). 2^nd^ divisions AHS evidence a significant reduction in the mitotic timing as a consequence of the rescue of cohesin function. **** = P<0.0001. **D-** Cohesin cleavage in the Wing disc of 3^rd^ *instar* larvae. Stills from live imaging showing single chromatids during mitosis. Dashed circles display epithelial cells from the wing disc pouch undergoing mitosis with loss of cohesion. **E-** Quantification of Cohesive states of divisions in the 3^rd^ *instar* larvae wing disc following RAD21-TEV cleavage and Rad21-WT rescue from 2h to 80h after heat-shock. Given the long cell cycle and the heterogeneous rate of division of cells in this tissue the presence of single sisters can still be observed up to 72h after the heat-shock. **F-** Quantification of mitotic timing and delay caused by Spindle Assembly Checkpoint activation after Rad21-TEV cleavage and Rad21-WT rescue in the 3^rd^ *instar* wing discs from 2 to 80hs AHS.In all panels n= number of cells. **** = P<0.0001; ** = P<0.01.

In contrast to the Nbs, in the epithelial cells of the wing disc, we observe the presence of single sisters and a mitotic delay even at 48hs AHS, despite the presence of high levels RAD21-WT (Figures 1A’’; 2D; 2E and 2F).These findings are consistent with the long cell cycle of the wing discs cells (27, 28). The high incidence of cells affected by reversible-cohesin cleavage is also consistent with a high frequency of cells in S/G2 in this tissue, estimated using the fly FUCCI system (29) (Figure S3B).To fully demonstrate the ability of our tool to induce aneuploidy in an acute manner in epithelial tissues we tested their regeneration capacity. In *Drosophila* epithelial cells, multiple cellular insults, including aneuploidy, can activate the Jun N-terminal kinase (JNK) signaling pathway, thus inducing the expression of pro-apoptotic genes and triggering the apoptotic cascade (9, 30). In agreement with these studies, 24hs AHS in the wing disc, Cleaved Caspase 3 (CC3) staining reveals a large population of dying cells thus reinforcing the notion that cell death is mostly a consequence of the induced chromosome segregation errors and the resulting aneuploidy (Figures S2A and S2A’). However, at 48hs AHS, the number of dying cells decreases significantly if cohesin activity is brought back, but not if the cohesion depletion by TEV is long-term (Figures S2A and S2A’). These results suggest that tissue recovery is limited and only possible if the mitotic disruption is restricted in time (or cell cycle), as achieved by our reversible genetic system. Although quantitative analysis was performed exclusively for wing discs epithelial cells and brain Nbs, analysis of other epithelial dividing tissues of the *Drosophila* larvae reveal a similar high incidence of single sisters 3hs AHS, implying that our system is able to induce a reversible-whole organism loss of cohesion (Figure S3A). We therefore conclude that our novel genetic tools is able to induce a single round of aberrant cell division, followed by quick rescue of mitotic fidelity, across the entire organism, leading to tissue-specific responses.

### Larvae challenged with aneuploidy during development hatch into impaired adults

To understand how the entire organism would respond to such high degree of induced chromosome segregation errors and consequent aneuploidy, we traced the larvae through development after cohesin cleavage. For comparative analysis, we monitored eclosion rates for organisms with the long-term TEV-protease cleavage system (inducing cohesin removal for >24hs) and our newly developed system with reversible inactivation of cohesin. Both systems represent a strong insult for all the dividing tissues of the larva; therefore, we expected them to be lethal in the pupa to adult transition. However, in contrast to several studies using chronic mitotic perturbations (10, 31) flies challenged with aneuploidy, using a reversible mitotic perturbation, ecloded into adult flies at high frequency, particularly if challenged up to 72hs after egg laying (AEL) (Figures 3A and Movie S3). Eclosion rates of adults were dependent on the developmental stage at which cohesin was reversibly cleaved (Figure 3A). Early induction of aneuploidy, at 48hs AEL, resulted in eclosion both with and without cohesin rescue. However, with 72hs AEL heat-shock, there was almost no eclosion if the RAD21 protein subunit was not brought back (Figure 3A). If the larvae were heat-shocked 96hs AEL, no cohesin rescue resulted in dead pupae, while cohesin rescue resulted in flies trying to escape the pupa, but unable to do so (“Head-out pupae”) (Figures 3A and 3B).These differences in developmental response to aneuploidy are likely due to increase of cell proliferation during larval development (10, 32) (Figure S1A). Regardless of the developmental stage, all flies that were able to eclode into adults after the aneuploidy challenge were completely unable to fly or move normally even when showing serviceable wings and appendages (Figure 3C’ and Movies S3 and S4).Consequently, these flies exhibited markedly shorter lifespans than their control counterparts (Figure 3C). Notably, even when aneuploidy is induced at earlier stages (48h AEL), thereby affecting fewer Nbs (Sup FIG1), neural tissue is not able to recover, with adults displaying “lethargic” behavior (Movie S5) while having an otherwise healthy adult morphology. This clearly exposes the different tissue sensitivities, showing that the developing brain is extremely sensitive to any level of aneuploidy during development.

**Figure 3.**
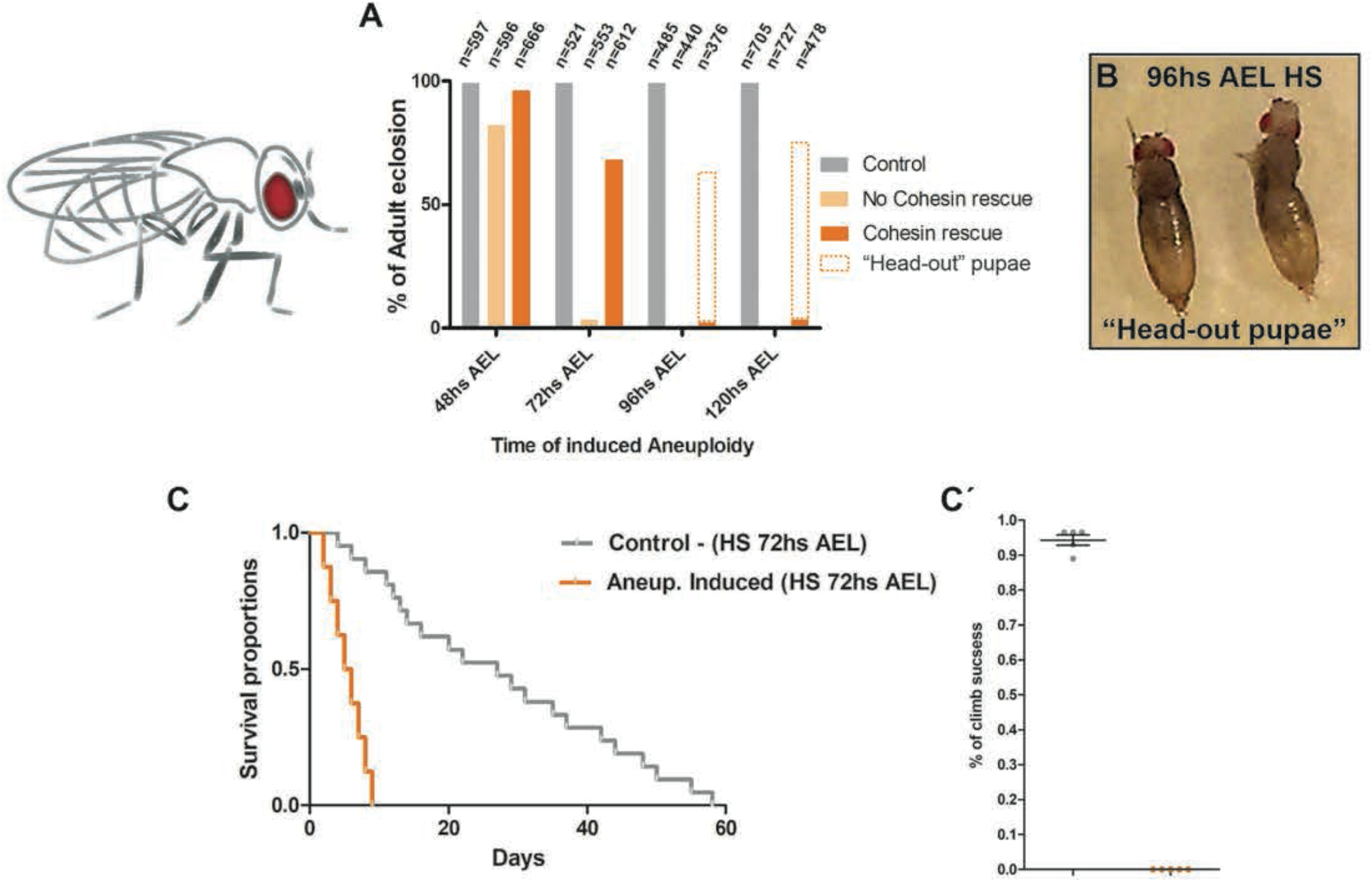
Larvae challenged with organism-wide aneuploidy hatch into adult flies with severe motor defects and reduced lifespan. **A-** Percentages of adult eclosions according to the stage of development at which reversible loss of cohesin (aneuploidy) was inducted (48, 72, 96 and 120hs after egg laying, AEL). n= number of flies.**B-** Picture showing “head out pupae” flies as a result of 96hs AEL induced aneuploidy **C to C′: C-** Kaplan-Meier survival curves showing fractional survival as a function of time. Ecloded flies with 72hs AEL induced aneuploidy showed reduced lifespan when compared with control flies (only heat-shocked). **C′-** Climbing assay comparing adult flies with 72hs induced aneuploidy and control flies. Percentage of climb success was plotted over the halfway point (10cm). Ecloded flies with 72hs AEL induced aneuploidy showed impaired motor behavior.

### Few cell cycles are sufficient to induce chromosomal instability in aneuploid Neuroblasts

We hypothesized that the severe motor defects in the newly hatched flies are a direct consequence of aneuploidy in the developing larva brain. Recently, it has been proposed that neural stem cells with unwanted karyotypes are eliminated (10, 31). To measure the number of neural stem cells over time after aneuploidy induction, we used the Nb marker Deadpan (DPN) to quantify all the nuclei with Nb morphology (Nb-like cells), defined based on their size, and located at the central brain area (CB) per lobe. The analysis indicates that there is a gradual decline in the Nbs number after the induction of aneuploidy from 12hs AHS onwards, but never a complete loss of the neural stem cell population (Figures 4A and 4A’). The slow kinetics and incomplete elimination of the stem cell population was quite surprising given the high levels of aneuploidy generated upon cohesin loss (~100%).

**Figure 4.**
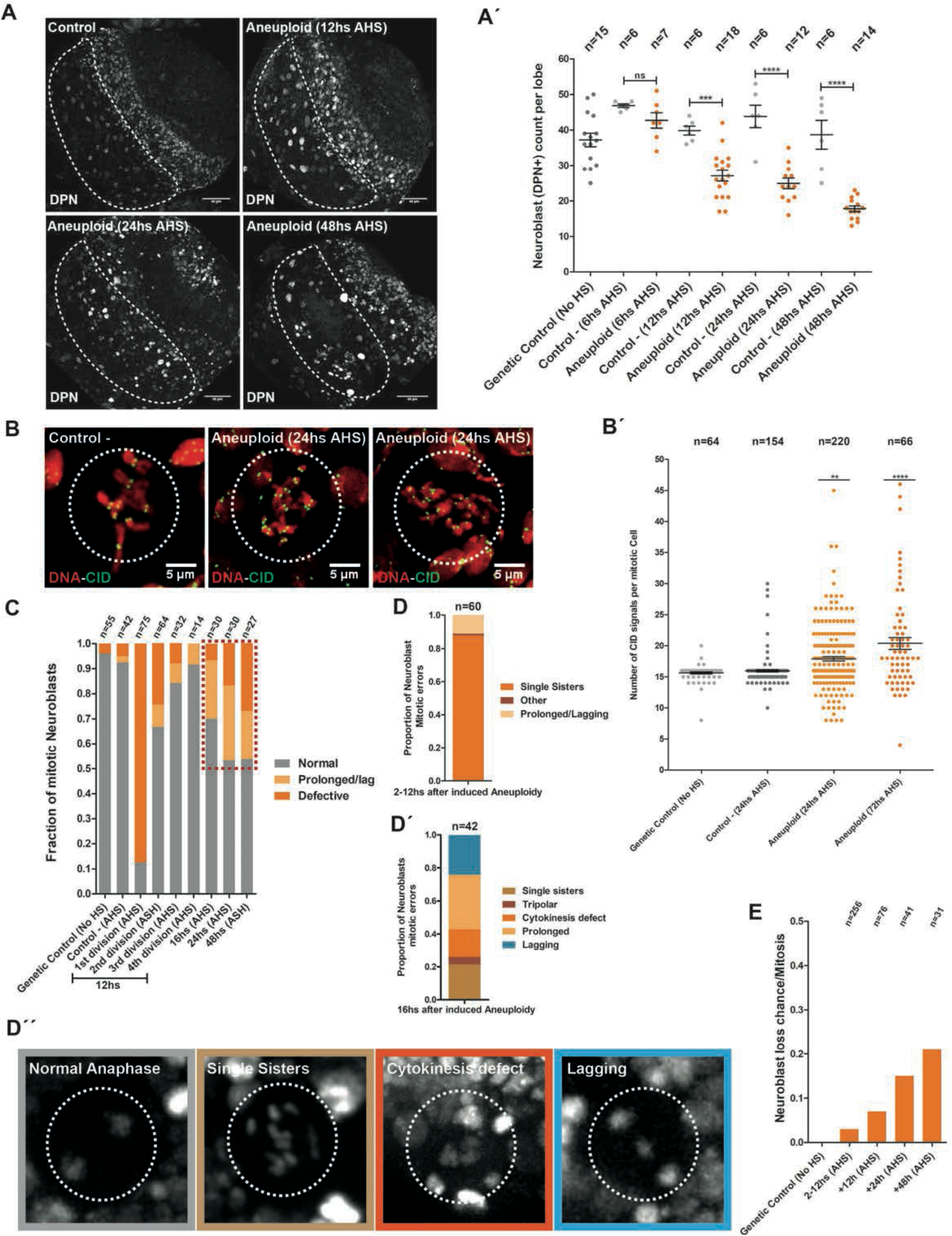
Aneuploidy results Neuroblast loss, chromosomal instability and accumulation of chromosomes. **A to AC’: A-** Nbs counts at the central brain (dashed shape) in 3^rd^ *instar* lobe brains assessed by immunofluorescence with the Nbs marker Deadpan (DPN). **A’-** Nbs were quantified based on the correlation between morphology and positive signal for DPN at 12, 24 and 48hs AHS. Reversible loss of cohesion and aneuploidy are followed by a reduction in Nbs numbers, but not a complete loss of the neural stem cell pool. n= number of lobe brains. *** = P<0.001; **** = P<0.0001; ns= no significant. Scale bar = 40μm. B to BC’: **B-** Chromosome counts were assessed by CID immunofluorescence (Centromere counts) in 3^rd^ *instar* Nbs arrested at metaphase with Colchicine (dashed circle) at 24 and 72hs AHS. B’-Aneuploid Nbs accumulate chromosomes through time. n= number of cells. **** = P<0.0001; ** = P<0.01. Scale bar = 5μm. **C-** Assessment of mitotic defects after aneuploidy induction from 2 to 48hs AHS. Chromosome instability arises shortly after aneuploidy induction (red dashed box). n= number of cells. **D to DC’C’: D and D’-** Profile of mitotic errors as a consequence of reversible cohesin depletion and consequent aneuploidy from 2 to 12hs AHS and 16 to 48hs AHS. D’’-Stills from live imaging documenting mitotic abnormalities in aneuploid Nbs (dashed circles). n= number of cells. **E-** Graph displaying the rate of catastrophic mitotic events which result in Nbs loss from 2 to 48hs AHS. Chromosomal instability can result in complete loss of Nbs morphology. n= number of cells.

Premature differentiation and apoptosis were suggested as the main mechanisms of aneuploid Nb elimination, reported in two recent studies (10, 31). However, after acute aneuploidy induction in the entire Nb population, we found a very low frequency of cells undergoing premature differentiation or cell death (Figure S4).As a proxy for premature differentiation events, we quantified Nb-like cells that had either lost the DPN marker or abnormally exhibit the differentiation marker Prospero (Pros) with or without co-expression of DPN (Figures S4A and S4A’, arrowheads and dashed circles).Pros is the key factor acting as a switch for the transition from stem cell self-renewal to terminal differentiation (33); therefore, this marker should not be present in Nbs. We observed that upon acute aneuploidy induction in the entire Nb population, there is a very low frequency of cells indicative of premature differentiation (Figure S4). These findings suggest that premature differentiation, although still taking place, is unlikely to be the major form of stem cell elimination. To estimate the levels of apoptosis, we also counted cells positive for cell death markers like CC3 and DCP1. We found a significant increase in CC3 positive cells in aneuploid brains (Figures S4A’; S4B’ and S4B’’) indicating that induction of apoptosis may also contribute to the elimination of aneuploid cells, as recently proposed (10). However, CC3 and DCP1 signals rarely correspond to Nb-like cells (For CC3 staining: Control: N=4 lobes Nbs/CC3=157/0 (0%) and Aneuploidy induced 24hs AHS: N=9 lobes Nbs/CC3=159/6 (3,7%); for DCP1 staining Control: N=5 lobes Nbs/CC3=201/0 (0%) and Aneuploidy induced 24hs AHS: N=10 lobes Nbs/CC3=191/5 (2,6%) (Figures S4B; S4B’ and S4B’’, arrowheads and dashed circles), suggesting that apoptosis may not be the major cause for NB elimination. Thus, loss of stem-cell identity and/or cell death are more likely potential consequences of genome randomization, rather than a specific mechanism controlling aneuploidy in the neural stem cell population (see discussion). Supporting this idea, inhibition of apoptosis by over-expression of the baculovirus protein P35 does not rescue Nbs number per brain lobe 24hs after induction of aneuploidy (Figure S4C).

To dissect the kinetics of the aneuploid response, we took advantage of the temporal resolution of our system allowing for the tracing of aneuploid fate in real time. We restricted our analysis to 3^rd^ *instar* wandering larvae as at this stage no new Nbs are generated from the neuro-epithelium (26). Induction of aneuploidy at this developmental stage, therefore, affects the entire Nbs population, which facilitates cell fate analysis. We observed a significant amount of Nbs proliferating for several days and displaying a tendency for chromosome accumulation over time (Figures 4B and 4B’). To analyze the number of chromosomes in each dividing Nbs we performed chromosome spreads and counted the number of centromeres per mitotic figure (each chromosome contains 2 centromere dots in mitosis). A single round of mitosis upon premature loss of sister chromatid cohesion should result in a maximum of 16 chromatids per Nbs, in the rare cases of complete asymmetric segregation (0,0013%). However, chromatid numbers can reach over 32 chromatids per cell, 24hs and 72h after loss of cohesion, at a much higher frequency (Figure 4B’). This analysis suggests that chromosome accumulation does not solely result from the initial perturbation.

To investigate this further, we characterized the mitotic fidelity of aneuploid Nbs. As described above, mitotic divisions that immediately follow the initial perturbation do not display significant mitotic errors and the low frequency of defects observed are cohesin-related (as expected from our experimental setup) (Figures 2B; 4C and 4D).

However, 16hs AHS, aneuploid cells start changing their behavior and a variety of mitotic defects appear, becoming more frequent over time (Figures 4C and 4D’). Detailed characterization of the mitotic defects arising 16hs after the induction of aneuploidy revealed that majority of them (~60%) are mild, consisting of either a prolonged metaphase or a lagging chromosome. However, the remaining ~40% consisted of cytokinesis defects, tri-polar spindles and sister chromatid cohesion defects, which are serious abnormalities that can drastically alter numerical ploidy (Figures 4D’ and 4D’’). This analysis further reveals that few hours are enough for the previously stable divisions of aneuploid karyotypes to become unstable, leading to further randomization of the genome. This chromosomal instability can also contribute to Nbs number decline, as catastrophic mitotic errors can result in complete loss of Nbs morphology and positioning (Figure 4E). All together, we conclude that upon transient cohesin loss and aneuploidy induction, neural stem cells exhibit a complex array of abnormalities as consequence of their karyotype diversification, such as loss of identity, cell death, or chromosomal instability, contributing to their gradual and loss over time.

### Karyotype restrictions in the proliferating aneuploid Neuroblast population

To test if there is a selection of specific karyotypes in the population of dividing aneuploid Nbs, we preformed Fluorescence *In Situ* Hybridization (FISH) analysis at 8hs and 24hs after aneuploidy was induced. To estimate the predicted frequency of specific karyotypes we first modeled the probability of each karyotype, assuming full random chromosome segregation in a single round followed by a second round of random segregation in ~20% of the cases (this was based on our experimental observations, see Figure 2B). FISH profiles were then compared with the statistical predictions (Figures 5A and 5B). The FISH profiles confirmed the propensity for chromosome accumulation over time (Figures 5C and 5D). Additionally, this analysis revealed that the karyotypes that can be tolerated by dividing Nbs are restricted to those containing at least one of the major three chromosomes, II III or X. The rate of complete loss of these chromosomes in the proliferating Nbs population was comparable to the control, and thus likely a consequence of experimental error of the FISH (Figures 5C’ and 5D’). We concluded that, although dividing aneuploid Nbs can persist in the tissue, their karyotypes have restrictions, as complete loss of any of the major three chromosomes prevents their proliferation in the developing brain. In contrast, other aneuploid combinations are compatible with continued proliferation, particularly when Nbs gain chromosomes.

**Figure 5.**
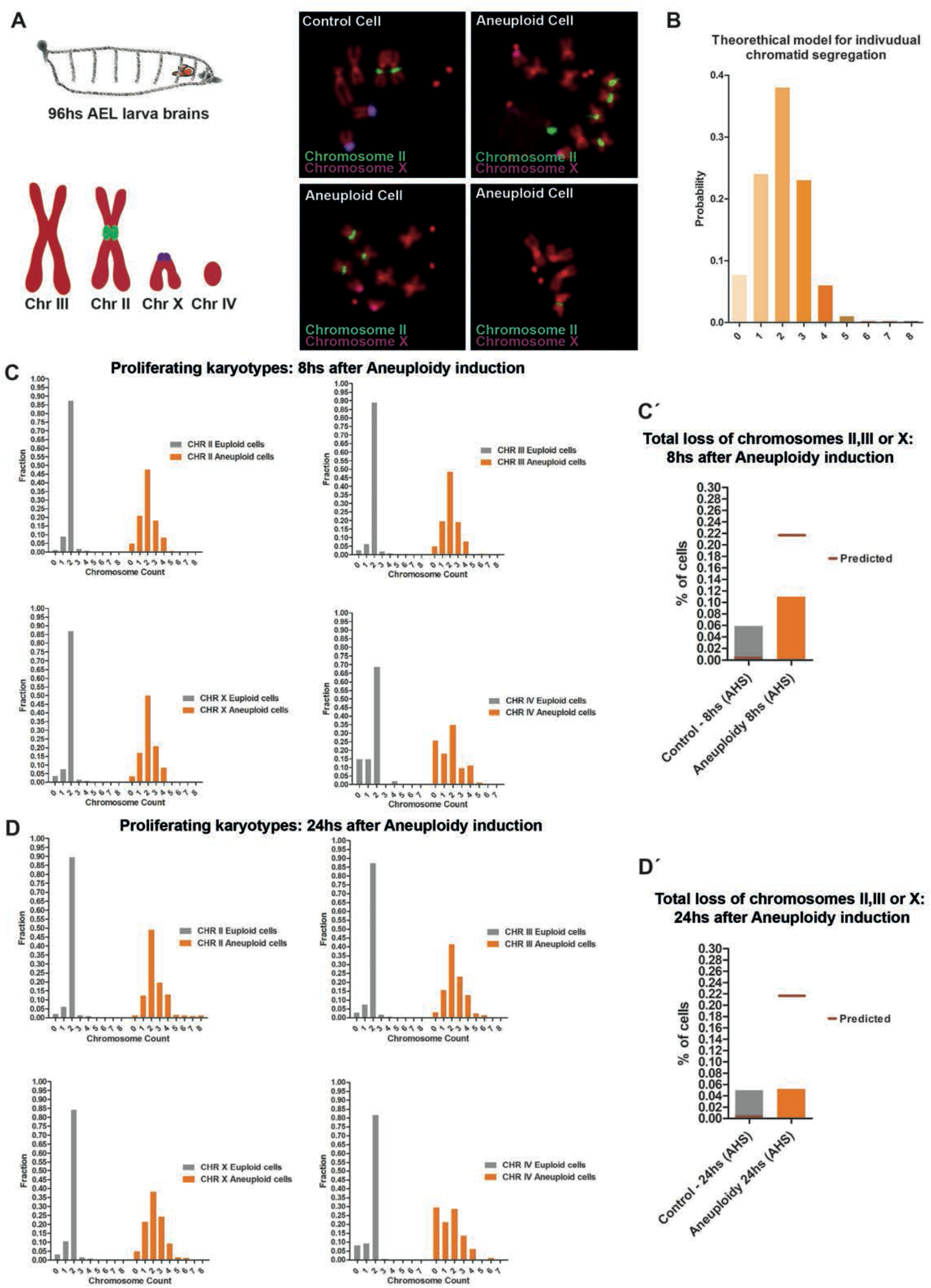
Karyotype restrictions in the proliferating aneuploid Nbs population. **A-** Panels of Fluorescent *in situ* Hybridization (FISH) of aneuploid Nbs and Control. Schematic of the FISH probe chromosome labeling. **B-** Theoretical segregation of sister chromatids of a given chromosome after cohesion loss, assuming segregation to be random (95% in the first mitosis, 20% in the second mitosis). We assumed that each chromatid segregates independently of all other chromatids and with equal probability to the mother and daughter cell. Given this, and for one cell division (mitosis) with diploid cells (4 chromatids for each chromosome) the distribution of the number of copies of each chromatid per cell is given by a Binomial distribution with 4 trials and probability of success of 0,5. C to **C′: C-** Frequency distribution of chromosome copy per Nbs, 8hs after aneuploidy induction. **C′-** Calculated theoretical loss rate for chromosome II, III or X, and the observed frequency of loss of any of these three chromosomes in the proliferating aneuploid Nbs (n>250 over eight brains analyzed per condition). **D to DC’: D-** Frequency distribution of chromosome copy per Nbs, 24hs after aneuploidy induction. D’-Calculated theoretical loss rate for chromosome II, III or X, and the observed frequency of loss of any of these three chromosomes in the proliferating aneuploid Nbs (n>250 over eight brains analyzed per condition).

### Aneuploidy elicits a stress response in the brain tissue

Our findings revealed that aneuploid cells are not promptly eliminated but instead continue to proliferate within certain karyotype restrictions. This should lead not only to the maintenance of aneuploid stem cells (due to Nb self-renewal) but also to the accumulation of differentiated aneuploid progeny (note that each Nb divides every ~2 hours (32)). We therefore tested how such increase in aneuploid cells within the tissue could affect cellular physiology and influence normal tissue development.

Several aneuploidy-associated stresses that include oxidative, metabolic, and proteotoxic stress are likely to alter cellular homeostasis (3), which ultimately lead to p53 activation and a p53-dependent cell-cycle arrest/senescence (34, 35). Interestingly, elevated levels of p53 have been observed in the Central Nervous System of Down syndrome patients (36). We decided to take advantage of our in vivo system to acutely induce aneuploidy to examine whether abnormal karyotypes trigger a stress response in the developing *Drosophila* brain and if so, what is the kinetics of such response. We assessed by immunohistochemistry the presence of P53 and the senescence marker Dacapo (DAP, a p21/p27 homologue (37)), after the loss of cohesin and consequent aneuploidy. We determined that both stress markers start to be evident at 12hs AHS in the tissue but only at 24hs AHS significant number of cells labeled with these markers are observed (Figures 6A; 6A’ and 6A’’). Furthermore, the large majority of the cells that appeared stress positive are not Nbs-like cells since the signal is limited to the small cells in the brain at that time (Figure 6A). Nbs-like cells stained with the stress markers are noticeable only at 48hs AHS (Figure 6A, arrowheads and dashed circles), suggesting that despite their aneuploid state, neural stem cells are delayed at displaying an evident stress-response. We confirmed this observation by quantifying the appearance of cells co-stained with the stress markers and the Nb marker DPN through time (Figure S5). We concluded that aneuploidy induction has a detectable effect in the entire brain population, triggering stress responses across different cell types. However, despite being promptly affected by aneuploidy induction, Nbs display a delay to this cellular insult and only mount a detectable response ~2 days after becoming aneuploid.

**Figure 6.**
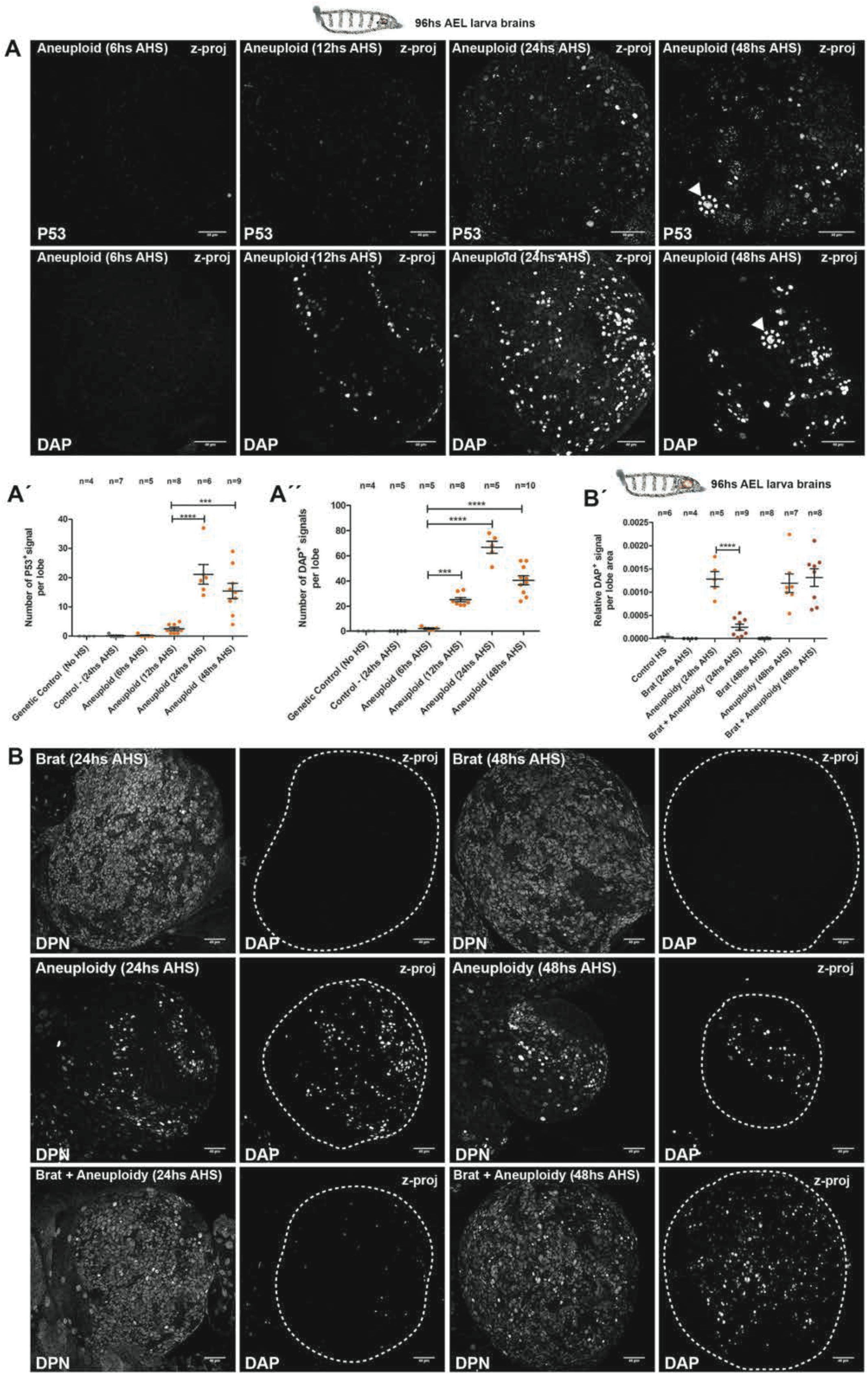
Aneuploidy-induced stress response is delayed in the neural tissue. **A to AC’C’: A-** Kinetics of the aneuploidy induced stress response at 6, 12, 24, and 48hs AHS; assessed by immunofluorescence of canonical markers P53 and Dacapo (DAP). Nbs show a delayed aneuploidy stress response at 48hs AHS (arrowheads with dashed circles). **AC’ and AC’’-** Counts of overall numbers of p53 and DAP signals per lobe, displaying a significant increase from 24hs AHS. ; *** = P<0.001; **** = P<0.0001. Scale bar = 40μm. z-proj (z projection). **B to BC’: B-** Pictures from fixed samples of 3^rd^ *instar* larvae lobe brains showing the immunofluorescence of the stress marker DAP and the Nbs marker (DPN) at 24 and 48hs after induction of aneuploidy. **B’-** Quantification of relative DAP positive signal per lobe area from 24 to 48hs AHS tissues. Brat mutant lobe brains showed a clear reduction in the presence of the aneuploidy induced stress marker DAP at 24hs AHS. n= number of lobe brains. **** = P<0.0001. Scale bar = 40μm. z-proj (z projection).

### Neural stemness delays aneuploidy stress response

The delayed stress response (i.e. ~48hs after induction of aneuploidy) in the neural stem cell pool may imply selective stem cell tolerance for the aneuploid condition when compared to the other cell types of the developing brain. To test this idea we took advantage of the *brat* mutant condition (38). In *brat* mutant larvae brains, each Nb divides into two daughter cells grow that retain Nbs properties, leading to the formation of a tumor-like neoplasm (39). We reasoned that cellular stemness confers tolerance to aneuploidy, the complete occupancy of the developing brain by Nbs-like cells observed in the *brat* mutant phenotype should be sufficient to prevent the stress response observed at 24hs AHS. To test this idea we combined our system for acute induction of aneuploidy with *brat* mutations to be able to induce aneuploidy in a *brat* mutant background and analyze the presence of stress markers at 24 and 48hs AHS. As predicted, DAP appearance was significantly delayed in aneuploid *brat* mutants when compared to aneuploid brains alone (Figures 6B and 6B’). The same result is observed for P53 staining (Figures S6A and S6A’). Note that the Nbs marker (DPN) stains almost all the cells in *brat* mutant brains, demonstrating the stem cell state of the entire tissue (Figures 6B and 6B’).This result suggests that the neural stem cell identity confers resistance to aneuploidy-associated stresses. Such delayed response has the drawback that enables continuous proliferation of aneuploid neural stem cells. The continued proliferation, in turn, allows for further brain growth despite presence of aneuploidy. Indeed, our results show that induced aneuploidy during larval development has no significant impact in the length of the brain and optic lobes in adult flies (Figure S7).

### Protecting only the developing brain from induced aneuploidy rescues the lifespan of ecloded flies

Upon aneuploidy challenge, we observed a striking difference across analyzed *Drosophila* tissues: whereas epithelial tissues like wing discs are able to regenerate from this insult (Figures S2A and S2A’), a significant fraction of neural stem cells continues proliferating and becomes highly chromosomally unstable (Figures 4A and 4C). These findings, together with the fact that most flies that survive the developmental aneuploidy induction show severe motor defects in otherwise healthy adult morphology, led us to hypothesize that the brain is the only limiting tissue in response to aneuploidy during fly development.

To test this hypothesis, we devised a system to selectively protect only the brain from cohesin removal and consequent aneuploidy. To achieve this, we complemented our reversible cohesin cleavage system with brain-specific expression of RAD21-WT throughout the course of the experiment (Figure 7A). In this way, TEV expression should lead to cohesion loss in all larval tissues that survive solely on RAD21-TEV at the time of heat shock. In contrast, neural stem cells should be resistant to this challenge, as they express both RAD21-TEV and RAD21-WT (Figure 7B). Neuroblast-specific expression of RAD21-WT was achieved by the use of *inscutable-Gal4 (insc-Gal4)* or *worniu-Gal4 (wor-Gal4)* drivers, to constitutively express *UAS-Rad21-wt-myc* in the developing brain (Figure 7B). As expected, constitutive presence of TEV-resistant RAD21 in the brain prevents any cohesion defects in 3^rd^ *instar* larvae Nbs (Figure 7C). To confirm that the rescue of sister chromatid cohesion occurs exclusively in the brain, we performed parallel characterization of the first mitotic division after the heat shock in the wing, derived from the same larvae. As anticipated, full cohesin cleavage was observed in all the dividing epithelial cells from the wing discs (Figure 7D). Notably, protecting only the brain from developmental aneuploidy fully rescued the severe motor defects of the ecloded flies from the 72hs AEL heat-shock, as demonstrated by mobility essays (Figures 7E’ and Movie S6).

**Figure 7.**
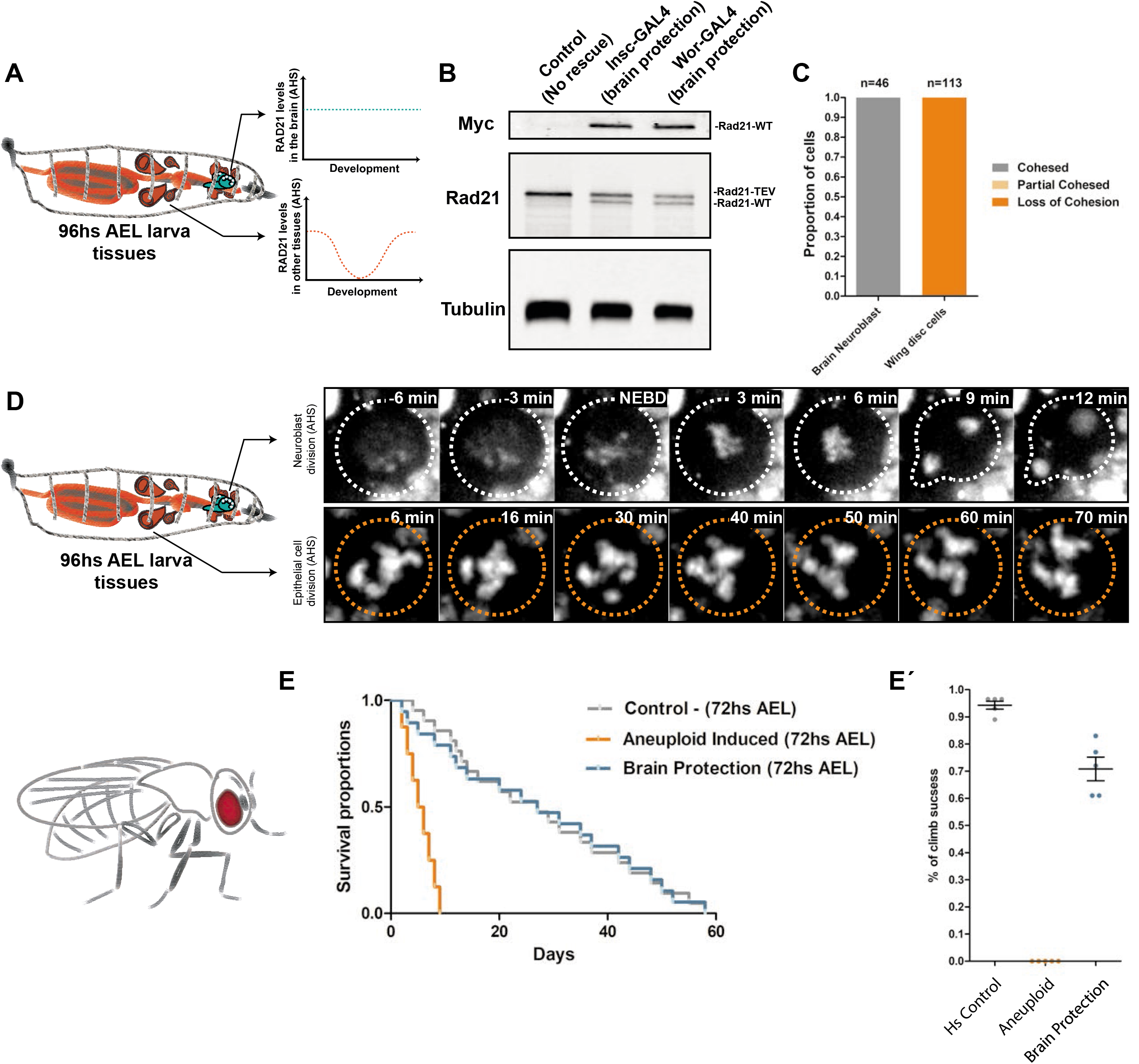
Protecting only the developing brain from induced aneuploidy rescues the adult lifespan. **A-** Graphic scheme depicting how the developing brain is protected from the loss of cohesion and induction of aneuploidy upon the constitutive expression of the RAD21-WT driven by Nbs specific Gal4 (*Insc-Gal4* or *Wor-Gal4)*. In contrast, the rest of the dividing tissues from the larva experience the acute inactivation of cohesin complex (by TEV cleavage of RAD21-TEV) after the heat-shock.**B-** Western blots showing the expression of both the cleavable (RAD21-TEV) and non-cleavable RAD21 (RAD21-WT) in 3^rd^ *instar* brains. *Insc-Gal4* and *Wor-Gal4* drivers result in expression of RAD21-WT in the 3^rd^ *instar* brain before the heat-shock. **C-** Quantification of Cohesive states of 3^rd^ *instar* larvae Nbs and epithelial cells from wing discs following heat-shock. *Insc-Gal4* protects the brain from cohesin loss, but has no effect in the wing disc. **D-** Live imaging cohesion profiles of Nbs and epithelial cells during mitotic divisions after the heat-shock. *Insc-Gal4* prevents mitotic delay caused by cohesion loss in the Nbs. E to EC’: **E-** Kaplan-Meier survival curves showing fractional survival as a function of time. Protection of the brain tissue from the induced aneuploidy rescues the adult lifespan of the ecloded flies with 72hs AEL heat-shock. **E’-** Climbing assay of adult flies. Percentage of climb success was plotted over the halfway point (10cm). Protection of the brain tissue from induced aneuploidy rescues motor defects of the ecloded flies with the 72hs AEL heat-shock.

Even more surprisingly, the brain protection was enough to rescue the lifespan of ~70% of the adult flies affected by organism-wide aneuploidy during development, demonstrating that the brain is indeed the most sensitive tissue when challenged with aneuploidy (Figure 7E).

## Discussion

### Acute disruption of mitotic fidelity enables tracing of aneuploidy *per se*

We developed a novel genetic tool in *Drosophila* to study aneuploidy *in vivo*. This tool enables the induction of a controlled pulse of aneuploidy, at the developmental stage of choice. The outcomes of using reversible perturbation are significantly different from the ones resulting from chronic disruption of mitotic fidelity. Whereas chronic mitotic perturbation is incompatible with organism viability, here we show a high survival rate upon controlled and acute organism-wide aneuploidy challenge. The long term survival after aneuploidy challenge coupled with the reversibility of the mitotic perturbation induced, overcomes one of the major limitations present in other metazoan models: we were able the study the kinetics response to aneuploidy across different tissues/developmental stages, focusing solely on the effects aneuploidy and without the confounding variable of the mitotic perturbation used to cause aneuploidy.

Cohesin loss and induction of aneuploidy is tolerated better by the organism if induced early in development, as observed by comparing the rates of eclosion. The developing larvae are progressively scaling mitotic machines, with each consecutive stage containing more divisions than the previous one (32). This implies that the heat-shock at the 1^st^ and 3^rd^ *instar* larvae are not the same, as they affect different number of dividing cells, thus generating different numbers of aneuploid progeny. Although, the more parsimonious explanation for aneuploidy tolerance in early development would be a quantitative one, it is also important to mention that a developmental delay is observed after aneuploidy induction (e.g. delayed pupariation stage). It is well known that delayed development allows the organism to adjust their growth programs after disturbances (40, 41). This induced delay is a development-stage dependent response, as some perturbations only appear to retard pupariation when induced at or before a certain stage in larval development as for example, beginning of the third *instar* (42-44).

### Chromosome mis-segregation in Neuroblasts leads to a complex array of karyotypes and cellular abnormalities

Neuroblasts have been used as a system to study aneuploidy response in previous studies (10, 31). These studies postulate two different but not mutually exclusive mechanisms of response to induced aneuploidy: premature differentiation (31) and cell death by apoptosis (10). We reasoned that if these are the major mechanisms of response to aneuploidy in neural stem cells, they should be detectable in high frequency after the aneuploidy induction by our acute approach. Contrary to that notion, after examined in detail the kinetics of the response, both premature differentiation and cell death were detected at low frequency even days after cells became aneuploid. It is important to note that the degree of aneuploidy in the Nbs upon cohesin loss should be around 98% due to the extensive genome shuffling prior to mitotic exit. Therefore, the finding that aneuploidy does not eliminate the entire Nb population, strongly argues against the existence of specific, active mechanisms controlling the integrity of the neural stem cell genome. The more plausible explanation is that the Nb elimination due to aneuploidy stems from a wide spectrum of abnormalities due to a randomized genome. Supporting this idea, it has been shown in yeast cell-to-cell variability in cell-cycle progression and robustness of multiple cellular processes even among cells harboring the same aneuploidies (45)

Examination of Nbs in real time after aneuploidy induction further revealed that aneuploidy is sufficient to induce chromosomal instability within a short time period (~16h). The appearance of obvious chromosomal instability, characterized by a wide range of mitotic defects, takes several cell cycles after cohesin has been restored, which strongly supports the notion that chromosomal instability is consequence of the abnormal karyotype and not the mitotic disruption initially applied. Overall, we observe a selection towards the accumulation of chromosomes, generating huge Nbs, which keep proliferating despite their increased ploidy (31). Thus, our *in vivo* detailed examination of the aneuploid Nbs (immediately after aneuploidy was induced) and kinetics of events (through several hours) clearly demonstrates that just a single round of chromosome mis-segregation in these cells is enough to originate a complex array of karyotypes which can lead to a variability of mitotic abnormalities.

### Neural stemness confers resistance to aneuploidy-associated stress response

During the last years, studies in tissue culture and yeast cells have collected solid evidence on how abnormal karyotypes can remarkably alter physiology of eukaryotic cells (reviewed in (46). They can lead to a different aneuploidy-associated responses that include oxidative, metabolic, and proteotoxic stress which likely contribute to p53 activation and cell senescence (34). However, our understanding about how aneuploidy-induced stress at the cellular level influences development of tissues is very limited.

Our time-course assessment of classical stress response markers (P53 and DAP) following chromosome mis-segregation in the brain tissue, clearly showed that aneuploidy response is not immediate and takes several hours for the cells to exhibit their up-regulation (12 to 24hs AHS). This delayed stress response is in agreement with recent observations in culture cells where it has been shown that chromosome mis-segregation did not lead to arrest in the following G1 in the vast majority of aneuploid daughter cells (47, 48).

Interestingly, our results highlight that cell identity determines the kinetics of this stress response. Aneuploidy response is specifically delayed in the neural stem cell pool (displayed mainly at ~48hs AHS) compared to the rest of the tissue, which exhibits it considerably earlier. Forcing self-renewal is sufficient to delay stress response in the entire tissue, suggesting that cellular stemness alone makes cells less sensitive to aneuploidy-induced stresses. Accordingly, unusual resistance to altered ploidy was observed in human and mouse embryonic stem cells (ESCs), mostly achieved by relaxing the cell cycle control and uncoupling the spindle checkpoint from apoptosis (49). The ability of neural stem cells to continue dividing despite the aneuploid karyotype dubbed them as aneuploidy “tolerant” (10). Yet, based on our findings it is clear that keeping these aneuploid cells is catastrophic for normal tissue architecture and development. Thus, aneuploidy may be “tolerated” better in Nbs, but the tissue as a whole is unable to be functional. In contrast, the “sensitivity” of epithelial cells enables the tissue to clean up and regrow properly.

### The developing brain restricts organism recovery after induced aneuploidy

Chromosomal aberrations have been long associated with neurological disorders (50). However, their impact on brain development and function remains complex and poorly understood, partially due to limitations of available experimental approaches. In almost all animal model systems used to study aneuploidy and its consequences until now, the organisms die prematurely due to the chronic disruption of mitotic fidelity to generate chromosome imbalance. Therefore, it is only possible to address the short term effect of aneuploidy in nervous system development, but not to understand the ultimate consequences for brain function. Our acute system reversibly affects chromosome segregation to induce just a pulse of aneuploidy, enabling the organism to recover from the insult and complete its development. The most noticeable phenotype observed in the adult was the severe motor and behavioral defects that clearly affect the lifespan of the flies, evidencing the sensitiveness of the nervous system to aneuploidy. Previous studies in *Drosophila* have shown that the mitotic disruption in larvae Nbs generates a reduction of their brain size (10, 31) reinforcing the idea about a link between aneuploidy and microcephaly. However, our results showed that induced acute aneuploidy has no significant impact in the size of the adult brain. These findings suggest that the continued proliferation of neural stem cells, caused by incomplete cell elimination and delayed aneuploidy-stress response, is sufficient to support the development of an apparently normal-sized organ. It is conceivable that the observed normal size reflects a sample selection, as this analysis was restricted to flies that survived the aneuploid challenge (~70%). Supporting this possibility, a screening performed to isolate anatomical brain mutants of *Drosophila* have shown that mutant strains showing altered brain shape and particularly small brains are very weak being mostly lethal at pupa stage (51). Despite unaltered shape and size of the adult brains, we reasoned that the neural circuits are likely impaired in those brains giving rise to the adult phenotype observed in all the surviving flies.

In accordance with the notion of the brain as the tissue most sensitive to aneuploidy, we show that preventing aneuploidy exclusively in the brain is sufficient to rescue all the behavioral defects previously observed. This brain protection not only rescued motor defects but also the lifespan of the flies ecloded upon 72hs AEL heat-shock, suggesting that neural tissue is the most ill-equipped to deal with aneuploidy during development and impose a significant cost for the organism. Several pathophysiological chromosomal disorders in humans including trisomy 21, trisomy 18, and trisomy 13, as well as the mosaic disorder MVA (mosaic variegated aneuploidy, characterized by the presence of a different number of chromosomes in some cells), are well-known to display intellectual disability (50), yet the impact of the aneuploid condition on brain development is still unclear (52, 53). Therefore, it becomes evident the necessity of future studies in different animal model systems based on an acute induction of aneuploidy to properly investigate its consequences for tissue development and homeostasis. These approaches could help to elucidate the molecular mechanisms underlying the physiological changes in stem/somatic cells generated by aneuploidy and its implications on tissue development and homeostasis.

## Methods

### Fly husbandry and genetics

Flies were raised using standard techniques at room temperature (20-22 °C). We established both chronic and the acute inactivation of cohesin complex by crossing the following genotypes: *w; hspr-nlsV5TEV; Rad21(ex^3^)/TM6B* with *w;; tubpr-Rad21(550-3TEV)-EGFP, Rad21(ex^15^), polyubiq-His-RFP*, and *w; hspr-nlsV5TEV; Rad21(ex^3^), hspr-Gal4, UAS-Rad21(wt)-myc/TM6B* with *w;; tubpr-Rad21(550-3TEV)-EGFP, Rad21(ex^15^), polyubiq-His-RFP*, respectively. The progeny was then heat shocked once at 37°C for 45min at the desired developmental stage. The correct genotype larvae were selected based on the absence of the “tubby” phenotype; the heat shocked “tubby” larvae were used as negative controls (control HS). As genetic control we used the same genotypes for the induction of aneuploidy but without performing the heat-shock.

To determine the proportion of adult eclosion, the crosses mentioned were raised in cages to monitor the time of egg collection. After 6hs collection, the plates were removed from the cages, the number of eggs counted and the plates were kept until larvae hatched. The plates were then heat-shocked at 37°C for 45min at different larvae developmental time (~48hs AEL, ~72hs AEL, ~96hs AEL and ~120hs AEL (± 6hs)) and placed in a new clean plastic cage. Once they reached pupae stage (“yellow body”) the pupae were gently removed with a wet brush and separated in “tubby” (control HS) and “no tubby” phenotype (condition). The different batches of pupae were placed over agar plates covered with two layers of absorbent paper to maintain the humidity and counted. The plates with the pupae were kept at room temperature until flies ecloded and the proportion of eclosion calculated.

To combine the induction of aneuploidy (acute cohesion inactivation) and the *brat* mutant genetic background we generated the following stocks: *w; brat^1^/CTG; Rad21(ex^3^), hspr-Gal4, UAS-Rad21(wt)-myc/TM6B* and w; *hspr-nlsV5TEV,brat^TS^/CTG; tubpr-Rad21(550-3TEV)-EGFP, Rad21(ex^15^), polyubiq-His-RFP*. These stocks were crossed and the progeny was heat shocked once at 37°C for 45min at the developmental stage desired and the genotype *w; brat^1^/hspr-nlsV5TEV, brat^TS^; Rad21(ex^3^), hspr-Gal4, UAS-Rad21(wt)-myc/tubpr-Rad21(550-3TEV)-EGFP, Rad21(ex^15^), polyubiq-His-RFP*, was selected at larva stage based on the absent of both, GFP signal and “tubby” phenotype.

To inhibit apoptosis we induced the over-expression of the baculovirus p35 in the context of the genetic background for acute inactivation of cohesin complex. To achieve this purpose, we generated the following stock *w; UAS-P35; tubpr-Rad21(550-3TEV)-EGFP 3, Rad21(ex^15^), polyubiq-His-RFP* to be crossed with *w; hspr-nlsV5TEV; Rad21(ex^3^), hspr-Gal4, UAS-Rad21(wt)-myc/TM6B*. The progeny was then heat-shocked once at 37°C for 45min at the developmental stage desired.

Finally, for the “brain rescue” experimental setup, we generated the following stocks: *w; insc-Gal4; tubpr-Rad21(550-3TEV)-EGFP, Rad21(ex^15^), polyubiqpr-His-RFP* and *w; wor-Gal4; tubpr-Rad21(550-3TEV)-EGFP, Rad21(ex^15^), polyubiqpr-His-RFP*. These stocks were crossed with the *w; hspr-nlsV5TEV; Rad21(ex^3^), hspr-Gal4, UAS-Rad21(wt)-myc/TM6B* stock. The crosses and the progeny were raised and treated as described above for the determination of the eclosion proportion.

Table with all stocks used in this study:

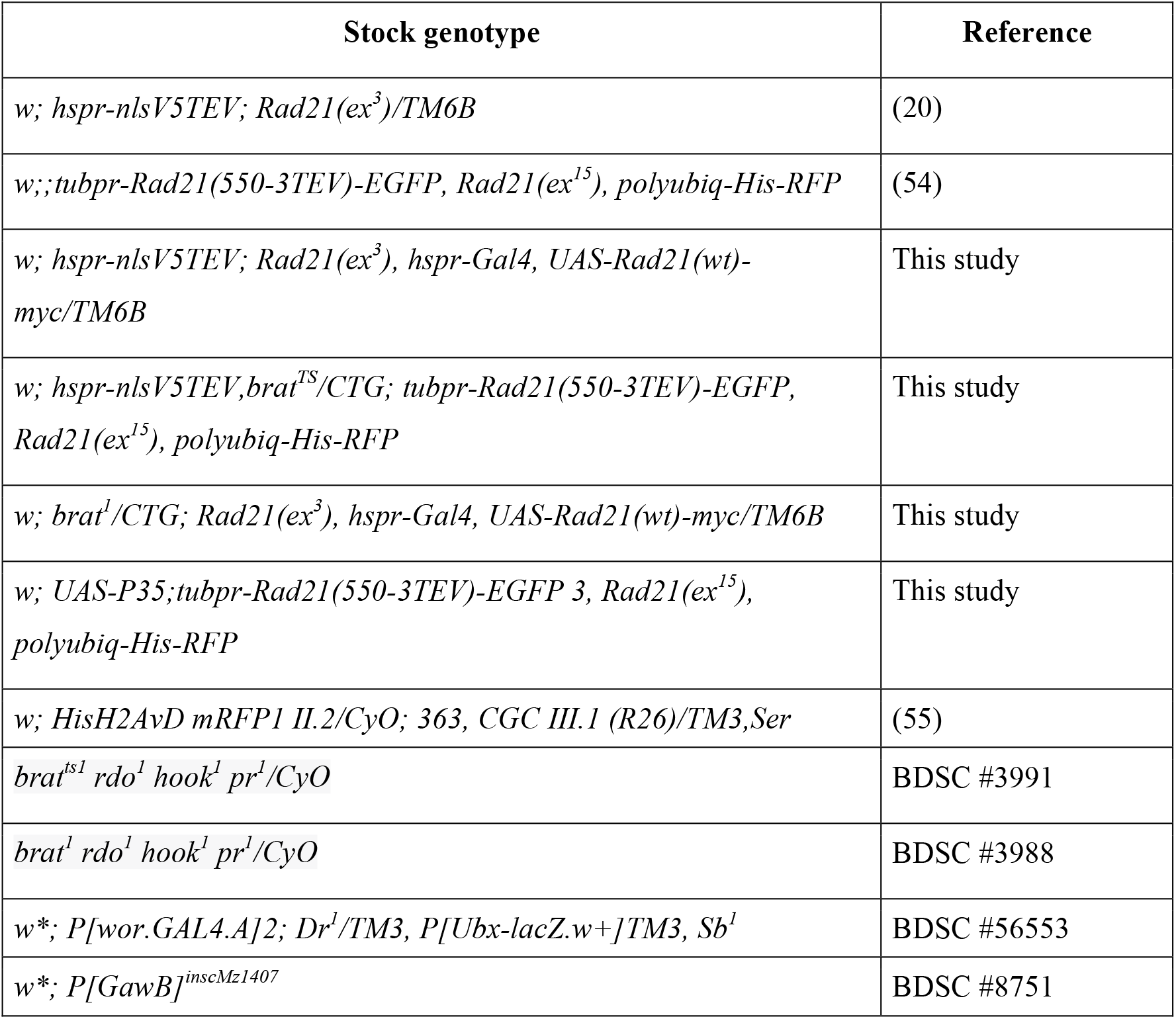

### Lifespan analysis

Lifespan was measured at room temperature according to standard protocols. In brief, newly ecloded animals (0 to 3 days) were collected (50 per genotype: “control”, “Aneuploidy” and “Aneuploidy + brain rescue”), and then placed in vials (up to 10 per vial), and transferred to fresh vials every two days. Survival was recorded for each vial. Due to the reduced mobility of the aneuploidy genotypes, we scored flies stacked in the food as death events in all the vials analyzed. We created survival curves with Prism 5.00 for Windows (GraphPad Software, San Diego, CA, USA) using the method of Kaplan and Meier.aq

### Climbing assay

For climbing assay flies were anesthetized with CO2, separated in groups of around twenty adults (3 replicas for each genotype) and allowed to recover for 2hs before beingsubjected to a climbing assay. Briefly, the groups of over twenty flies were placed in an empty climbing vial and then tapped down to the bottom. They were allowed to climb past the halfway point from the bottom of the vial for 30 seconds (10cm). The number of flies above the 10 cm mark was recorded as a percentage of flies able to climb.

### Histology

Briefly, flies were anesthetized with CO2 and then were placed gently in agarose blocks to immobilize them and prevent any damage to the head or eyes. The agarose blocks with the flies were immersed in Carnoy fixative overnight, at 4°C. The next day the Carnoy solution was removed and three 70% ethanol washes were performed. Immediately after, the flies were decapitated and the heads were oriented one by one in melted 2% agarose to guarantee similar orientation of the tissue sections. Agarose blocks were then processed, embedded, the whole head was sectioned into 5um-thick sequential sections and stained with Hematoxylin & Eosin. The histology was performed in the Histopathology unit at Instituto Gulbenkian de Ciência and the slides were analyzed by a pathologist with a DMLB2 microscope (Leica). Images were acquired with a DFC320 camera (Leica) and NanoZoomer-SQ Digital slide scanner (Hamamatsu).

### Live-cell imaging

Larvae 3^rd^ *instar* brains were dissected in Schneider medium supplemented with 10% FBS and intact brains were mounted on a glass-bottom dish (MakTek), covered with an oxygen-permeable membrane (YSI membrane kit), and sealed with Voltalef oil 10S (VWR). This procedure allowed long-term imaging of brains for periods up to 10 hours.

For imaging of imaginal discs and early *instar* larvae brains, tissues were dissected in Schneider medium with 10% FBS. Dissected discs were placed and oriented in a 200μl drop of medium at the bottom of a glass-bottom dish (MakTek).

Live imaging was performed on a spinning disc confocal using imaged on a Revolution XD microscope (Andor, UK) equipped with immersion a 60x glycerol-immersion 1.30 NA objective (Leica Microsystems) and a iXon Ultra 888 1024*1024 EMCCD (Andor, UK). 25-35 Z-series optical sections were acquired 0.5-1 μm apart.

### Brain spreads and Immunofluorescence

For brain spreads and immunofluorescence, 3^rd^ *instar* larvae brains were dissected in PBS, incubated with 100 μM colchicine for one hour, hypotonic shocked in 0.5% sodium citrate for 2-3 minutes, and fixed on a 5 μl drop of fixative (3.7% formaldehyde, 0.1% Triton-X100 in PBS) placed on top of a siliconized coverslip. After 30 seconds, the brains were squashed between the coverslip and a slide, allowed to fix for an additional 1 min and then placed in liquid nitrogen. Slides were further extracted with 0.1% Triton-X100 in PBS for 10 min, and used for immunofluorescence following standard protocols. Primary antibodies were rat anti-CID (gift from Claudio E. Sunkel) used at 1:2000, Cleaved *Drosophila* Dcp-1 (Asp216) Antibody (1:300) # 1679578S (Cell Signaling Technology), Cleaved Caspase-3 (Asp175) Antibody #9661 (1:300) (Cell Signaling Technology), Anti-Deadpan antibody #ab195173 (1:1500) (Abcam). Secondary antibodies conjugated with fluorescent dyes from Alexa series (Invitrogen) were used according to the manufacturer’s instructions.

Third *instar* wing imaginal disc fixation and staining, as well as immunofluorescence of whole brains was performed using standard procedures (Lee and Treisman, 2001). Briefly, third *instar* larvae wing disc tissue (still attached to the larva body) was fixed on ice for 30 min. The fixative consisted of 4% formaldehyde (Polysciences) in 1X PEM buffer solution. Following were washed by gentle agitation three times for 20 min in PBS-T (1x PBS + 0.1% Triton X-100). Primary antibodies incubation was performed overnight at 4 °C in PBS-T supplemented with 1% BSA and 1% donkey serum. The following day, the tissues were washed again and incubated for 2h at room temperature with the appropriate secondary antibodies diluted in PBS-T solution. Finally, after the wash of secondary antibodies, wing discs were mounted in Vectashield (Vector Laboratories). Fluorescence images were acquired with a ×40 HCX PL APO CS oil immersion objective (numerical aperture: 1.25–0.75) on a Leica SP5 confocal microscope.

### Fluorescence *In Situ* Hybridization

Brains from 3^rd^ instar larvae were dissected in PBS, incubated with 100 μM colchicine for one hour, and transferred to 0.5% sodium citrate solution for 3-4 minutes. Then, the brains were transferred to a fixative containing 11:11:2 Methanol: Acetic Acid: MQ Water, for 30 seconds before being placed in a droplet of 45% Acetic acid for 2 minutes, squashed and transferred to liquid Nitrogen. Then, the coverslip was removed and the slide incubated in absolute ethanol for 10 min at −20 °C (Freezer incubation). The slides were air dried at 4 °C. (20 minutes).The slides were dehydrated at room temperature in 70%, 90% and absolute ethanol for 3 minutes, prior to DNA denaturation in 70% formamide-2xSCC solution for 2 minutes at 70 °C. This is done on the thermomixer set at 70 °C with a formamide solution heated to70 °.Then, the slides were transferred to cold 70% Ethanol (−20 °C) and dehydrated at room temperature in 90% and absolute ethanol for 3 min. FISH probes were denatured in the hybridization buffer at 92C for 3 min. Hybridization was done over-night at 37 °C using 30 ul of FISH hybridization buffer/probe mix per slide. Hybridization buffer: 20% dextran sulfate in 2x SCCT/50% Formamide/0,5mg/ml Salmon sperm DNA. Then, slides were washed 3 × 5 min in 50% formamide-2xSCC at 42 °C and 3 × 5 min in 0.1xSCC at 60 °C. These steps are done on the thermomixer, with the solutions previously heated to desired temperatures. Finally, the slides are washed in PBS, and mounted in Vecta shield with DAPI. The probes were used in the final concentration of 70Nm in hybridization buffer. Probes used were: Chr_X (359 bp satellite DNA) A546-GGGATCGTTAGCACTGGTAATTAGCTGC, and Ch_3 (dodeca satellite DNA) Cy5-ACGGGACCAGTACGG DNA probes, Chr_2 A488-(AACAC).

### Western-blot

To analyze RAD21 protein amounts, *Drosophila* tissues were dissected in PBS and homogenized with a pestle in Sample buffer. Samples were centrifuged, and boiled for 5 minutes in 2x Sample Buffer. Samples were loaded on a 13 % SDS-gel for electrophoresis and and then transferred to nitrocellulose membranes. Western-blot analysis was performed according to standard protocols using the following antibodies: anti-α-tubulin (1:50.000, DM1A, Sigma-Aldrich Cat# T9026), guinea pig anti-Rad21 (Heidmann et al., 2004) and V5 Tag Mouse Monoclonal Antibody (Novex^®^).

### Image analysis

Imaging analysis was performed using FIJI software (Schindelin et al., 2012). For z-projections slices were stacked into maximum intensity (10 frames, 2μm each). Some pictures were rotated and/or flipped to orient them in the same way.

### Statistical analysis

Statistical analysis and graphic representations were performed using Prism 5.00 for Windows (GraphPad Software, San Diego, CA, USA). Unpaired t test or one-way ANOVA (using the Bonferroni’s multiple comparison) were applied depending the measurements analyzed in the corresponding experiment. Sample size details are included in the respective plotted graphs.

## Acknowledgments

We thank S. Heidmann and the Bloomington Stock Center for fly strains, the Advance Imaging Unit and Fly Facility, Histopathology of Instituto Gulbenkian de Ciencia for technical assistance, and C. Homem, F. Janody and M. Bettencourt-Dias, and all the members of the RAO laboratory for discussions and comments. MM was supported by a Fundação para a Ciência e Tecnologia, FCT, fellowship (SFRH /BD/52438/2013). This work was supported by the following grants awarded to RAO: FCT Investigator grant (IF/00851/2012/CP0185/CT0004), EMBO Installation Grant (IG2778) and European Research Council Starting Grant (ERC-2014-STG-638917).

---

**Supplementary Figure 1.**
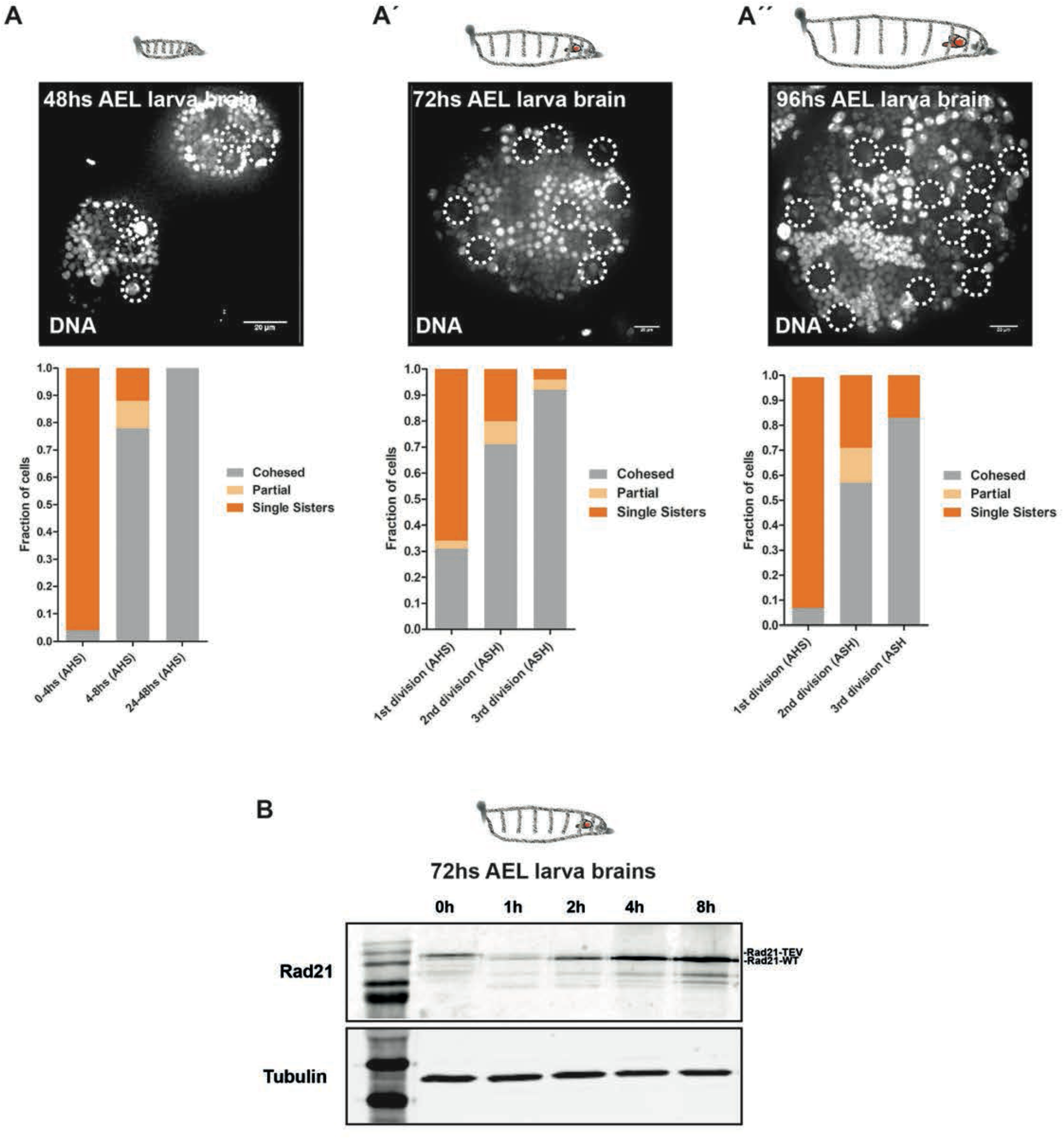
Heat-shock treatment induces brain aneuploidy at all stages of development. **A to AC’’-** Stills from live imaging of lobe brains at different larvae stages (48, 72 and 96hs AEL); dashed circles are highlighting the Nbs in the lobes (N>3 brains per condition). The number of dividing Nbs increase with larvae development. Cohesive state of Nbs after the loss of cohesion and subsequent rescue in 48, 72 and 96hs AEL larvae were plotted.b **B-** Western blot of RAD21 cleavage and rescue dynamics in 72hs AEL larvae brains (over 10 western blots were performed to validate the system).

**Supplementary Figure2.**
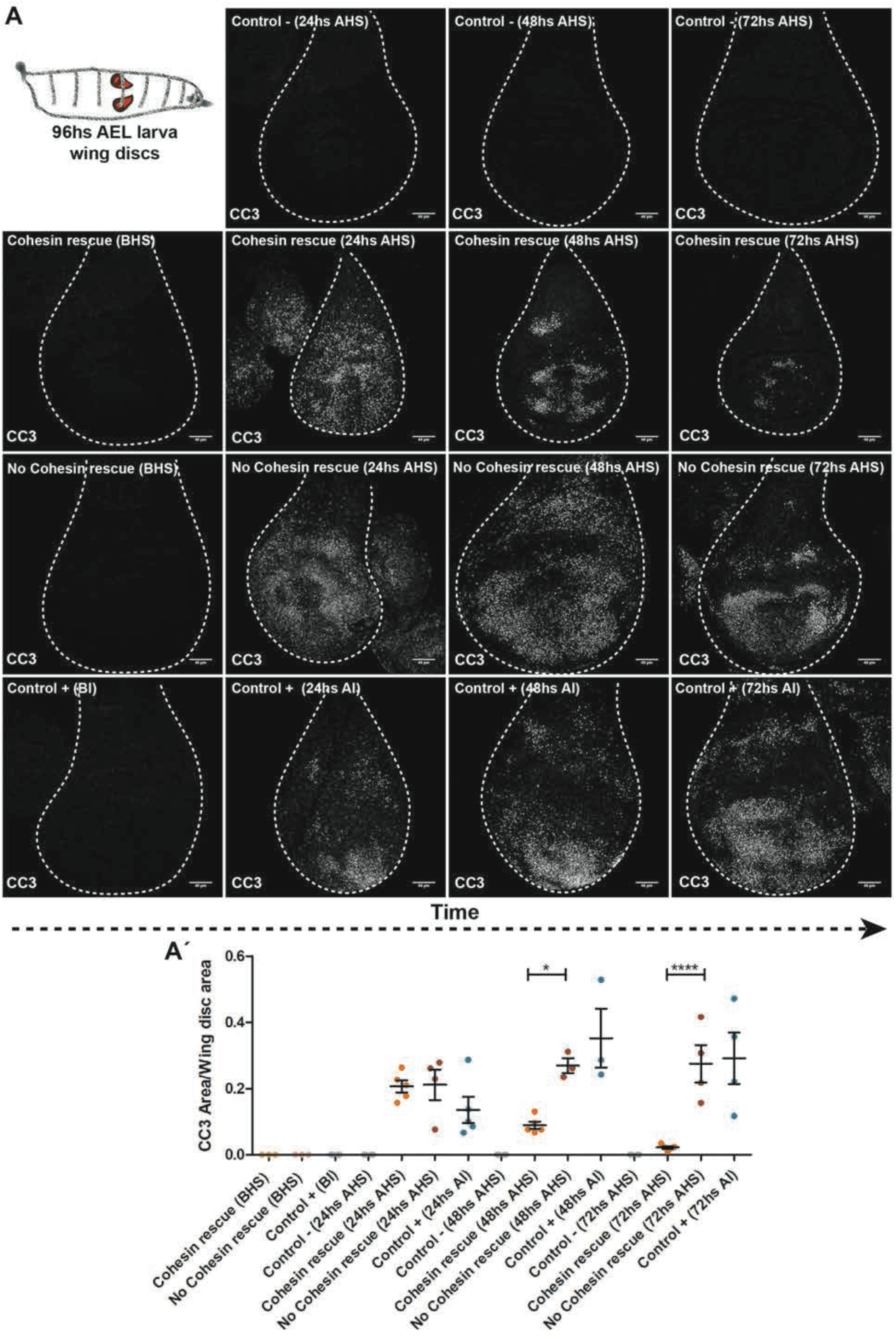
Epithelial tissues recover from high levels of aneuploidy by cell death and compensatory proliferation. **A to AC’: A-** Reversible cohesin cleavage results in apoptosis in the 3^rd^ *instar* wing discs (dashed shapes depict the wing disc areas). The amount of apoptosis per disc was measured by area of cleaved caspase 3 (CC3) immunofluorescence at 24, 48 and 72hs AHS. **A’-** Rescue of cohesin function significantly reduced the amount of apoptosis within 48hs after of the heat shock. In contrast, chronic inactivation of cohesin complex (No cohesin rescue) displayed high levels of apoptosis through time. Control-(Control HS); Control+ (Irradiation: 4,000 rads); BHS (Before Heat-Shock, genetic control); AHS (After Heat-Shock, condition); BI (Before Irradiation); AI (After Irradiation). * = P<0.05; **** = P<0.0001. Scale bar = 40μm. z-proj (z projection).

**Supplementary Figure 3.**
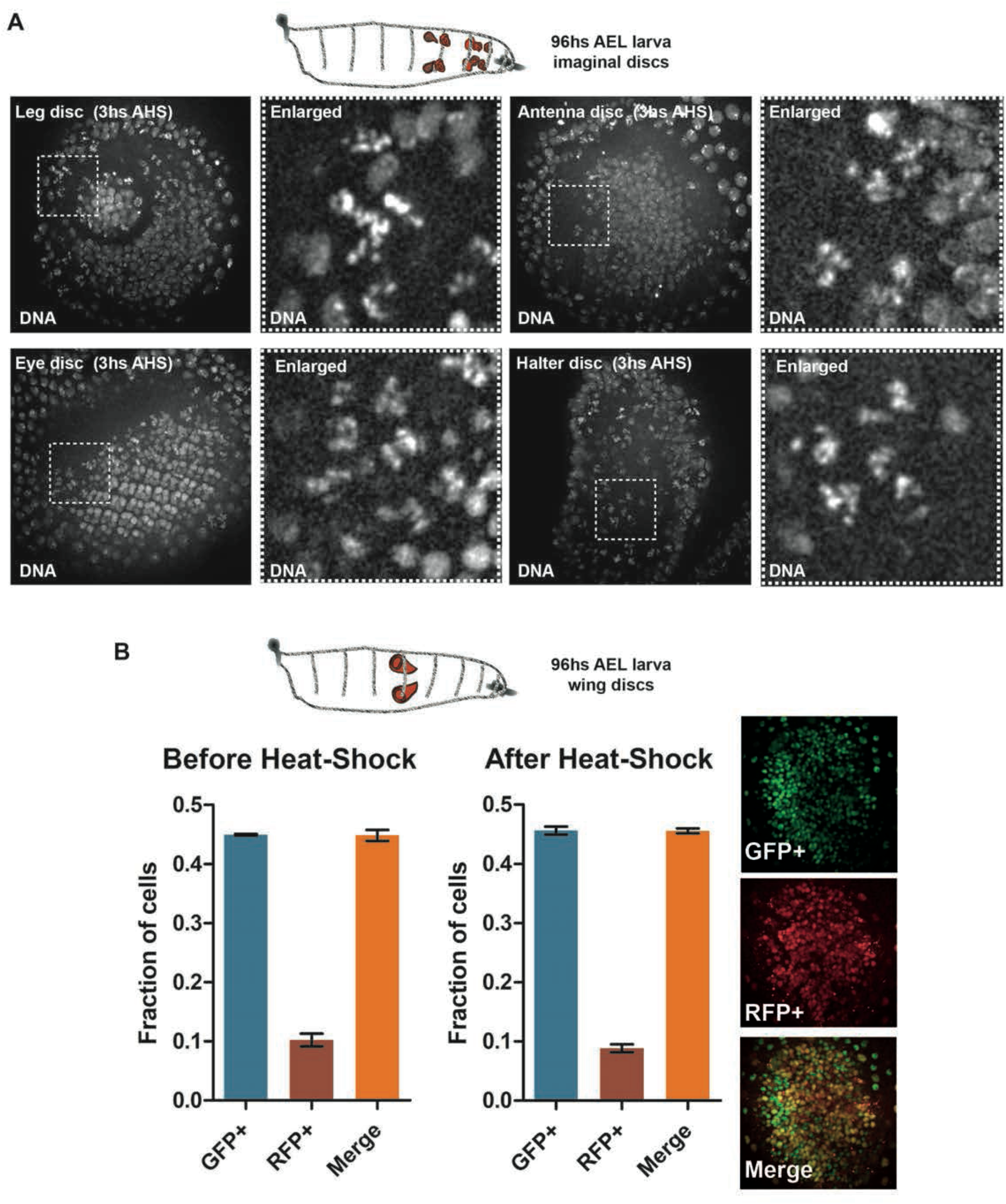
RAD21 cleavage and rescue induces loss of cohesion in all examined dividing tissues. **A-** Stills from live imaging of leg, eye, antennae and halter 3^rd^ *instar* imaginal discs after induction of RAD21 cleavage. Dashed squares display epithelial cells from the imaginal discs undergoing mitosis with loss of cohesion (see enlarged picture). **B-** The cell cycle profile evaluation of the 3^rd^ *instar* control wing disc with or without the heat shock, using the FLY-FUCCI system. The high incidence of cells affected by reversible cohesin cleavage is consistent with a high frequency of cells in G2/M in this tissue (see Merge). GFP: G1 cells; RFP: S phase cells; Merge: G2/M Cells (n>500, at least three wing discs analyzed)

**Supplementary Figure 4.**
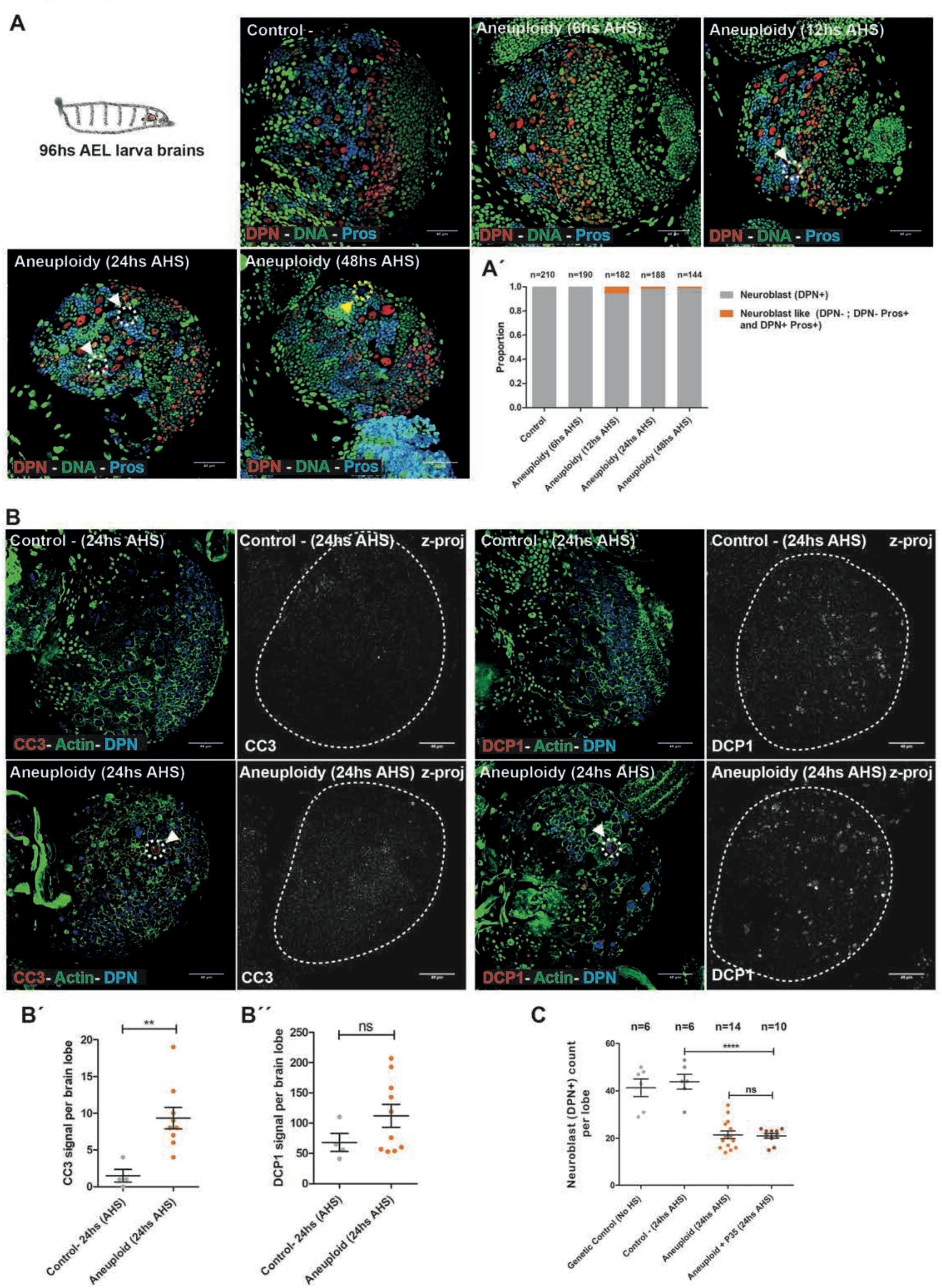
Aneuploidy results in low frequency of stem identity loss and cell death in Neuroblasts. **A to AC’: A-** Pictures from fixed samples of 3^rd^ *instar* larvae lobe brains stained with Deadpan (DPN), Prospero (Pros) and Histone RFP (DNA). Induction of aneuploidy results in the loss of stem cell identity measured by the absent of Deadpan (stem cell marker, white arrowhead with dashed circle), appearance of Prospero (differentiation marker, yellow arrowhead with dashed circle) or both markers together in cell nucleus with “Nbs shape like”. **A’-** Percentage of loss of stem cell identity in the neural stem cell pool at different time points after the induction of aneuploidy. These events are observed at very low frequency. n= number of Nbs like cells. Scale bar = 40μm **B to BC’C’: B-** Pictures from fixed samples of 3^rd^ *instar* larvae lobe brains stained with Deadpan (DPN), Cleaved Caspase 3 (CC3, death marker), DCP1 (death marker) and rhodamine phalloidin (Actin). Induction of aneuploidy results in cell death measured by the presence of CC3 or DCP1 signals (white arrowheads with dashed circles) in cells with “Nbs shape like”. **BC’ and BC’C’-** Quantification of cell death signals CC3 and DCP1 per larvae brain lobes at 24hs AHS. The presence of positive signal for the cell death markers in Nbs shape like cells is very low. ** = P<0.01. ns= not significant. Scale bar = 40μm. z-proj (z projection) **C-** Quantification of Nbs at the central brain in 3^rd^ *instar* lobe brains assessed by immunofluorescence with the Nbs marker DPN. Inhibition of apoptosis by over-expression of baculovirus P35 does not rescue Nbs number after 24hs induced aneuploidy. n= number of lobe brains. **** = P<0.0001. ns= not significant.

**Supplementary Figure 5.**
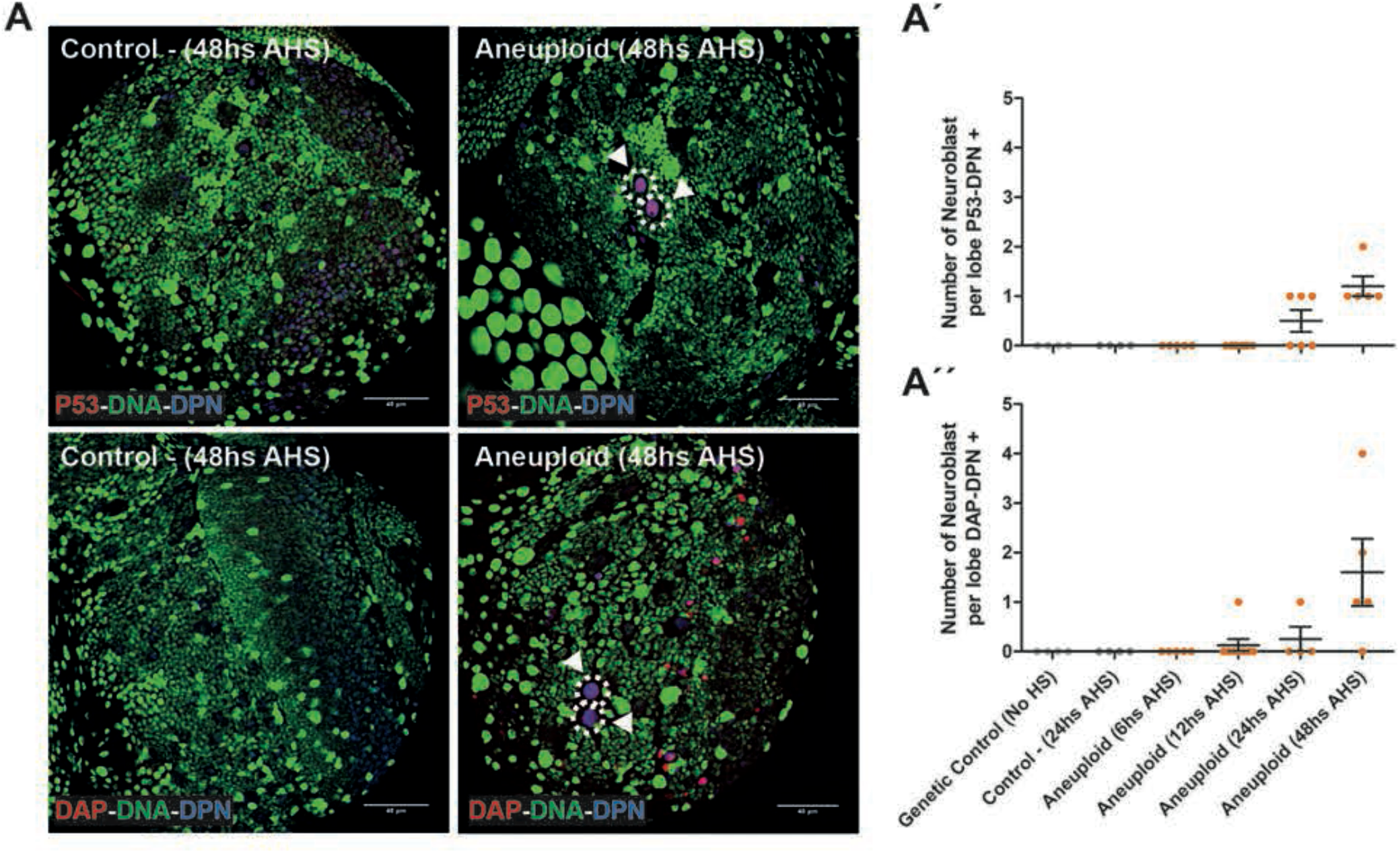
Aneuploidy induced stress response is particularly delayed in the Neuroblasts. **A to AC’C’:** Pictures from fixed samples of 3^rd^ *instar* larvae lobe brains showing the immunofluorescence of canonical stress response markers P53 and Dacapo (DAP) together with the Nbs marker (DPN) at 48hs AHS. Nbs display a delayed aneuploidy stress response at 48hs AHS (arrowheads with dashed circles). **AC’ and A’’-** Quantification of the kinetics of the aneuploidy induced stress response at 6, 12, 24, and 48hs AHS in Nbs (DPN+). Scale bar = 40μm.

**Supplementary Figure 6.**
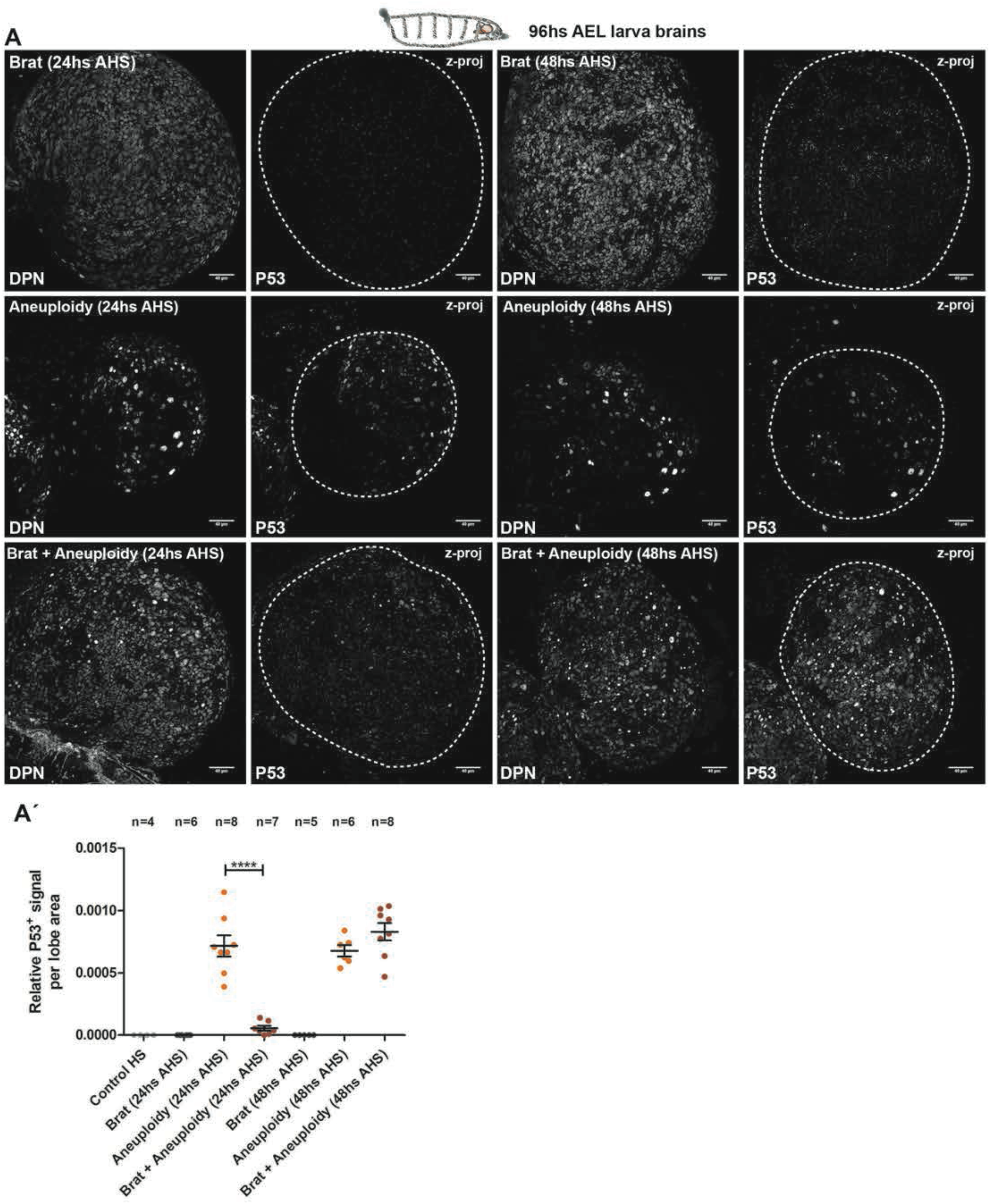
Aneuploidy induced P53 accumulation is delayed in the neural stem cell pool. **A to AC’: A-** Brat mutant lobe brains showed a clear reduction in the presence of the aneuploidy induced stress marker P53 at 24hs AHS. **A’-** Quantification of relative P53 positive signal per lobe area from 24 to 48hs AHS. n= number of lobe brains. **** = P<0.0001. Scale bar = 40μm. z-proj (z projection).

**Supplementary Figure 7.**
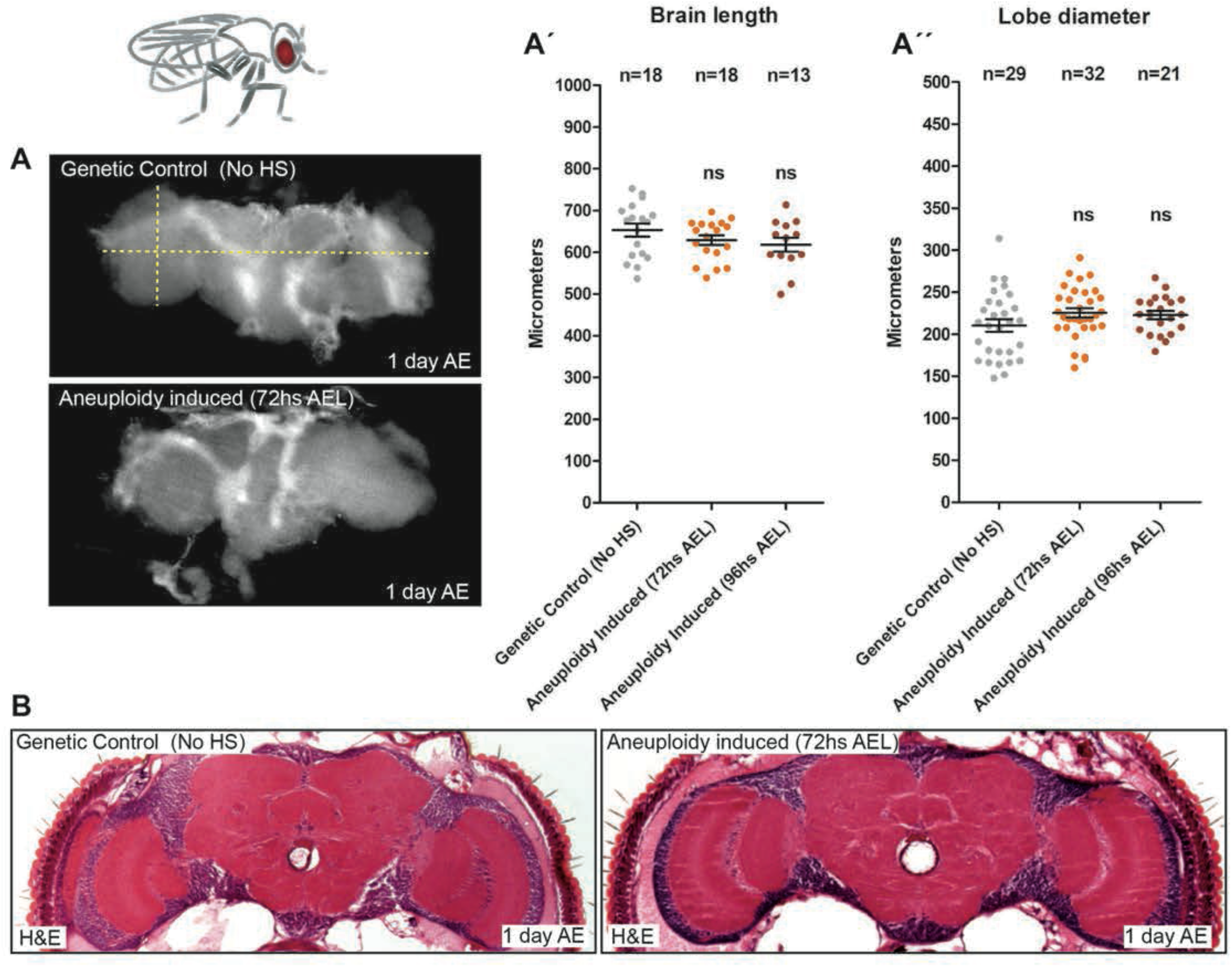
Adult brains do not show any significant alteration in size after acute induction of aneuploidy during development. **A to A’’: A-** Dissected brains of adult flies from a control and a developmental aneuploidy-induced (72hs AEL heat-shock) organisms. **A’ and A’’-** Quantifications of lobe diameter and brain length in control and developmental aneuploidy-induced (72 and 96hs AEL heat-shock) adult flies showed no significant differences. n= number brains. ns= no significant **B-** Histology analysis of brains from control and aneuploidy-induced during development (72hs AEL heat-shock) adult flies, one day after eclosion. Frontal sections at approximately midbrain showed no signal of neurodegenerative process (vacuolization). H&E= Hematoxylin and Eosin.

## #Supplementary Movies

**Supplementary Movie 1: Demonstration of cohesin cleavage and rescue system in a single Neuroblast after the heat shock**.

Movie depicting live imaging of the first and second mitosis after the heat chock, in a single 3^rd^ instar larvae Neuroblast; Imaging of HisH2AvD mRFP1 in 3 minute timeframes. First division after the heat shock results in complete cohesin loss, while the second provides a full cohesin rescue.

**Supplementary Movie 2: Demonstration of cohesin cleavage and rescue in the entire larval brain lobe after the heat shock**.

Movie depicting live imaging of the 3^rd^ instar larvae brain lobe; Imaging of HisH2AvD mRFP1 in 3 minute timeframes. First divisions after the heat shock result in complete cohesin loss and genome shuffling, consecutive divisions result in cohesin rescue.

**Supplementary Movie 3: Cohesin loss and rescue results in eclosion of adults with motion defects, despite healthy wing and appendage morphology**.

Movie depicting live imaging of the adult eclosed after the aneuploidy challenge at the age of 72hs after egg laying.

**Supplementary Movie 4: A Climbing assay comparing heat shock controls and flies challenged with aneuploidy during development**.

**Supplementary Movie 5: 48hs AEL Heat-Shock results in eclosion of adults with lethargic behavior despite healthy wing and appendage morphology**.

Movie depicting live imaging of the adult eclosed after the aneuploidy challenge at the age of 48hs after egg laying

**Supplementary Movie 6: Climbing assay comparing brain-rescued flies and flies challenged with aneuploidy during development**.

## References

1. Ambartsumyan G, Clark AT. Aneuploidy and early human embryo development. Hum Mol Genet. 2008;17(R1):R10–5.

2. Holland AJ, Cleveland DW. Boveri revisited: chromosomal instability, aneuploidy and tumorigenesis. Nat Rev Mol Cell Biol. 2009;10(7):478–87.

3. Santaguida S, Amon A. Short- and long-term effects of chromosome mis-segregation and aneuploidy. Nat Rev Mol Cell Biol. 2015;16(8):473–85.

4. Sheltzer JM, Ko JH, Replogle JM, Habibe Burgos NC, Chung ES, Meehl CM, et al. Single-chromosome Gains Commonly Function as Tumor Suppressors. Cancer Cell. 2017;31(2):240–55.

5. Gordon DJ, Resio B, Pellman D. Causes and consequences of aneuploidy in cancer. Nat Rev Genet. 2012;13(3):189–203.

6. Lara-Gonzalez P, Westhorpe FG, Taylor SS. The spindle assembly checkpoint. Curr Biol. 2012;22(22):R966–80.

7. Knouse KA, Wu J, Whittaker CA, Amon A. Single cell sequencing reveals low levels of aneuploidy across mammalian tissues. Proc Natl Acad Sci U S A. 2014;111(37):13409–14.

8. Pfau SJ, Silberman RE, Knouse KA, Amon A. Aneuploidy impairs hematopoietic stem cell fitness and is selected against in regenerating tissues in vivo. Genes Dev. 2016;30(12):1395–408.

9. Dekanty A, Barrio L, Muzzopappa M, Auer H, Milan M. Aneuploidy-induced delaminating cells drive tumorigenesis in Drosophila epithelia. Proc Natl Acad Sci U S A. 2012;109(50):20549–54.

10. Poulton JS, Cuningham JC, Peifer M. Centrosome and spindle assembly checkpoint loss leads to neural apoptosis and reduced brain size. J Cell Biol. 2017;216(5):1255–65.

11. Ly P, Cleveland DW. Interrogating cell division errors using random and chromosome-specific missegregation approaches. Cell Cycle. 2017:1–7.

12. Hassold T, Hunt P. To err (meiotically) is human: the genesis of human aneuploidy. Nat Rev Genet. 2001;2(4):280–91.

13. Guacci V, Yamamoto A, Strunnikov A, Kingsbury J, Hogan E, Meluh P, et al. Structure and function of chromosomes in mitosis of budding yeast. Cold Spring Harb Symp Quant Biol. 1993;58:677–85.

14. Michaelis C, Ciosk R, Nasmyth K. Cohesins: chromosomal proteins that prevent premature separation of sister chromatids. Cell. 1997;91(1):35–45.

15. Mirkovic M, Oliveira RA. Centromeric Cohesin: Molecular Glue and Much More. Prog Mol Subcell Biol. 2017;56:485–513.

16. Nasmyth K, Haering CH. Cohesin: its roles and mechanisms. Annu Rev Genet. 2009;43:525–58.

17. Haering CH, Farcas AM, Arumugam P, Metson J, Nasmyth K. The cohesin ring concatenates sister DNA molecules. Nature. 2008;454(7202):297–301.

18. Ivanov D, Nasmyth K. A topological interaction between cohesin rings and a circular minichromosome. Cell. 2005;122(6):849–60.

19. Mirkovic M, Hutter LH, Novak B, Oliveira RA. Premature Sister Chromatid Separation Is Poorly Detected by the Spindle Assembly Checkpoint as a Result of System-Level Feedback. Cell Rep. 2015;13(3):470–8.

20. Pauli A, Althoff F, Oliveira RA, Heidmann S, Schuldiner O, Lehner CF, et al. Cell-type-specific TEV protease cleavage reveals cohesin functions in Drosophila neurons. Dev Cell. 2008;14(2):239–51.

21. Gerlich D, Koch B, Dupeux F, Peters JM, Ellenberg J. Live-cell imaging reveals a stable cohesin-chromatin interaction after but not before DNA replication. Curr Biol. 2006;16(15):1571–8.

22. van Ruiten MS, Rowland BD. SMC Complexes: Universal DNA Looping Machines with Distinct Regulators. Trends Genet. 2018;34(6):477–87.

23. Eichinger CS, Kurze A, Oliveira RA, Nasmyth K. Disengaging the Smc3/kleisin interface releases cohesin from Drosophila chromosomes during interphase and mitosis. EMBO J. 2013;32(5):656–65.

24. Rao SSP, Huang SC, Glenn St Hilaire B, Engreitz JM, Perez EM, Kieffer-Kwon KR, et al. Cohesin Loss Eliminates All Loop Domains. Cell. 2017;171(2):305–20 e24.

25. Urbach R, Technau GM, Breidbach O. Spatial and temporal pattern of neuroblasts, proliferation, and Engrailed expression during early brain development in Tenebrio molitor L. (Coleoptera). Arthropod Struct Dev. 2003;32(1):125–40.

26. Homem CC, Knoblich JA. Drosophila neuroblasts: a model for stem cell biology. Developmentp. 2012;139(23):4297–310.

27. Milan M, Campuzano S, Garcia-Bellido A. Cell cycling and patterned cell proliferation in the wing primordium of Drosophila. Proc Natl Acad Sci U S A. 1996;93(2):640–5.

28. Neufeld TP, de la Cruz AF, Johnston LA, Edgar BA. Coordination of growth and cell division in the Drosophila wing. Cell. 1998;93(7):1183–93.

29. Zielke N, Korzelius J, van Straaten M, Bender K, Schuhknecht GF, Dutta D, et al. Fly-FUCCI: A versatile tool for studying cell proliferation in complex tissues. Cell Rep. 2014;7(2):588–98.

30. Milan M, Clemente-Ruiz M, Dekanty A, Muzzopappa M. Aneuploidy and tumorigenesis in Drosophila. Semin Cell Dev Biol. 2014;28:110–5.

31. Gogendeau D, Siudeja K, Gambarotto D, Pennetier C, Bardin AJ, Basto R. Aneuploidy causes premature differentiation of neural and intestinal stem cells. Nat Commun. 2015;6:8894.

32. Ito K, Hotta Y. Proliferation pattern of postembryonic neuroblasts in the brain of Drosophila melanogaster. Dev Biol. 1992;149(1):134–48.

33. Choksi SP, Southall TD, Bossing T, Edoff K, de Wit E, Fischer BE, et al. Prospero acts as a binary switch between self-renewal and differentiation in Drosophila neural stem cells. Dev Cell. 2006;11(6):775–89.

34. Kruiswijk F, Labuschagne CF, Vousden KH. p53 in survival, death and metabolic health: a lifeguard with a licence to kill. Nat Rev Mol Cell Biol. 2015;16(7):393–405.

35. Thompson SL, Compton DA. Proliferation of aneuploid human cells is limited by a p53-dependent mechanism. J Cell Biol. 2010;188(3):369–81.

36. Liao JM, Zhou X, Zhang Y, Lu H. MiR-1246: a new link of the p53 family with cancer and Down syndrome. Cell Cycle. 2012;11(14):2624–30.

37. Lane ME, Sauer K, Wallace K, Jan YN, Lehner CF, Vaessin H. Dacapo, a cyclin-dependent kinase inhibitor, stops cell proliferation during Drosophila development. Cell. 1996;87(7):1225–35.

38. Arama E, Dickman D, Kimchie Z, Shearn A, Lev Z. Mutations in the beta-propeller domain of the Drosophila brain tumor (brat) protein induce neoplasm in the larval brain. Oncogene. 2000;19(33):3706–16.

39. Betschinger J, Mechtler K, Knoblich JA. Asymmetric segregation of the tumor suppressor brat regulates self-renewal in Drosophila neural stem cells. Cell. 2006;124(6):1241–53.

40. Gontijo AM, Garelli A. The biology and evolution of the Dilp8-Lgr3 pathway: A relaxin-like pathway coupling tissue growth and developmental timing control. Mech Dev. 2018.

41. Hackney JF, Cherbas P. Injury response checkpoint and developmental timing in insects. Fly (Austin). 2014;8(4):226–31.

42. Garelli A, Gontijo AM, Miguela V, Caparros E, Dominguez M. Imaginal discs secrete insulin-like peptide 8 to mediate plasticity of growth and maturation. Science. 2012;336(6081):579–82.

43. Halme A, Cheng M, Hariharan IK. Retinoids regulate a developmental checkpoint for tissue regeneration in Drosophila. Curr Biol. 2010;20(5):458–63.

44. Simpson P, Berreur P, Berreur-Bonnenfant J. The initiation of pupariation in Drosophila: dependence on growth of the imaginal discs. J Embryol Exp Morphol. 1980;57:155–65.

45. Beach RR, Ricci-Tam C, Brennan CM, Moomau CA, Hsu PH, Hua B, et al. Aneuploidy Causes Non-genetic Individuality. Cell. 2017;169(2):229–42 e21.

46. Santaguida S, Vasile E, White E, Amon A. Aneuploidy-induced cellular stresses limit autophagic degradation. Genes Dev. 2015;29(19):2010–21.

47. Santaguida S, Richardson A, Iyer DR, M'Saad O, Zasadil L, Knouse KA, et al. Chromosome Mis-segregation Generates Cell-Cycle-Arrested Cells with Complex Karyotypes that Are Eliminated by the Immune System. Dev Cell. 2017;41(6):638–51 e5.

48. Soto M, Raaijmakers JA, Bakker B, Spierings DCJ, Lansdorp PM, Foijer F, et al. p53 Prohibits Propagation of Chromosome Segregation Errors that Produce Structural Aneuploidies. Cell Rep. 2017;19(12):2423–31.

49. Mantel C, Guo Y, Lee MR, Kim MK, Han MK, Shibayama H, et al. Checkpoint-apoptosis uncoupling in human and mouse embryonic stem cells: a source of karyotpic instability. Blood. 2007;109(10):4518–27.

50. Bushman DM, Chun J. The genomically mosaic brain: aneuploidy and more in neural diversity and disease. Semin Cell Dev Biol. 2013;24(4):357–69.

51. Heisenberg B. Isolation of Anatomical Brain Mutants of Drosophila by Histological Means. Z Naturforsch 34 c, 143–147 (1979) 1979.

52. Oromendia AB, Amon A. Aneuploidy: implications for protein homeostasis and disease. Dis Model Mech. 2014;7(1):15–20.

53. Ricke RM, van Deursen JM. Aneuploidy in health, disease, and aging. J Cell Biol. 2013;201(1):11–21.

54. Oliveira RA, Kotadia S, Tavares A, Mirkovic M, Bowlin K, Eichinger CS, et al. Centromere-independent accumulation of cohesin at ectopic heterochromatin sites induces chromosome stretching during anaphase. PLoS Biol. 2014;12(10):e1001962.

55. Oliveira RA, Hamilton RS, Pauli A, Davis I, Nasmyth K. Cohesin cleavage and Cdk inhibition trigger formation of daughter nuclei. Nat Cell Biol. 2010;12(2):185–92.

